# The Hippo pathway controls myofibril assembly and muscle fiber growth by regulating sarcomeric gene expression

**DOI:** 10.1101/2020.10.08.330951

**Authors:** Aynur Kaya-Çopur, Fabio Marchiano, Marco Y. Hein, Daniel Alpern, Julie Russeil, Nuno Miguel Luis, Matthias Mann, Bart Deplancke, Bianca H. Habermann, Frank Schnorrer

**Affiliations:** Aix Marseille University, CNRS, IBDM, Turing Center for Living Systems, 13288 Marseille, France; Max Planck Institute of Biochemistry, 82152 Martinsried, Germany; Institute of Bioengineering, School of Life Sciences, École Polytechnique Fédérale de Lausanne (EPFL), 1015 Lausanne, Switzerland

**Keywords:** muscle, sarcomere, myofibril, Hippo, yorkie, *Drosophila*, growth, Dlg5, STRIPAK

## Abstract

Skeletal muscles are composed of gigantic cells called muscle fibers, packed with force-producing myofibrils. During development the size of individual muscle fibers must dramatically enlarge to match with skeletal growth. How muscle growth is coordinated with growth of the contractile apparatus is not understood. Here, we use the large *Drosophila* flight muscles to mechanistically decipher how muscle fiber growth is controlled. We find that regulated activity of core members of the Hippo pathway is required to support flight muscle growth. Interestingly, we identify Dlg5 and Slmap as regulators of the STRIPAK phosphatase, which negatively regulates Hippo to enable post-mitotic muscle growth. Mechanistically, we show that the Hippo pathway controls timing and levels of sarcomeric gene expression during development and thus regulates the key components that physically mediate muscle growth. Since Dlg5, STRIPAK and the Hippo pathway are conserved a similar mechanism may contribute to muscle or cardiomyocyte growth in humans.

## Introduction

Mammalian skeletal muscles are built from gigantic cells called muscle fibers, up to several centimetres long, that mechanically link distant skeletal elements. Muscle forces are produced by highly regular molecular arrays of actin, myosin and titin filaments called sarcomeres. Each sarcomere has a length of about three micrometres in relaxed human skeletal muscles (Ehler and Gautel, 2008; Llewellyn et al., 2008; Regev et al., 2011). Thus, hundreds of sarcomeres need to assemble into long chains called myofibrils in order to generate force across the entire muscle fiber (Lemke and Schnorrer, 2017). Large muscle fibers contain many parallel myofibrils, which are laterally aligned to a cross-striated pattern to effectively power animal locomotion (Gautel, 2008; Schiaffino et al., 2013). How muscle fibers grow to these enormous sizes and how their growth is coordinated with the assembly and growth of the individual myofibrils within the muscle is a challenging biological problem that is not well understood.

Muscle fibers are built during animal development. Initially, many small myoblasts fuse to myotubes, whose long-ends then mechanically connect to tendon cells (Kim et al., 2015; Schnorrer and Dickson, 2004). This enables the build-up of mechanical tension within myotubes, which consecutively triggers myofibril assembly and the transition of myotubes to early myofibers (Weitkunat et al., 2017; 2014). Following myofibril assembly, the immature myofibrils mature and build functional sarcomeres. To do so each myofibril grows in length and diameter and thereby supports the extensive muscle fiber growth during embryonic and postembryonic development (González-Morales et al., 2019; Orfanos et al., 2015; Reedy and Beall, 1993; Sanger et al., 2017; Sparrow and Schöck, 2009). For the correct developmental sequence of myofibril morphogenesis, the protein concentrations of the various sarcomeric components need to be precisely regulated (Orfanos and Sparrow, 2013; Schönbauer et al., 2011). This is particularly prominent in mammalian muscle fibers, in which sarcomeric proteins transcriptionally switch isoforms from embryonic to neonatal and finally adult isoforms (Schiaffino, 2018; Schiaffino et al., 2015). In *Drosophila* indirect flight muscles, transcription of sarcomeric protein coding genes starts just before myofibril assembly and is then strongly boosted during myofibril maturation, when myofibrils grow in length and width (González-Morales et al., 2019; Shwartz et al., 2016; Spletter et al., 2018). Concomitantly with the growth of the myofibrils, also the mitochondria grow in size (Avellaneda et al., 2020). How this precise transcriptional control is achieved and coordinated with muscle fiber growth is unclear.

One central pathway controlling organ size during development and tumorigenesis is the Hippo pathway, which regulates the activity of the growth promoting transcriptional coactivator Yorkie (Yki, YAP and TAZ in mammals) (Pan, 2010; Zanconato et al., 2019). The core of the pathway is composed of a kinase cascade with Hippo (Hpo; Mst1 and Mst2 in mammals) phosphorylating the downstream kinase Warts (Wts; Lats1 and Lats2 in mammals) (Udan et al., 2003; Wu et al., 2003). Phosphorylated Wts is active and in turn phosphorylates Yki (Huang et al., 2005), leading to the cytoplasmic retention of Phospho-Yki by 14-3-3 proteins (Dong et al., 2007; Oh and Irvine, 2008; Ren et al., 2010). When the pathway is not active unphosphorylated Yki enters into the nucleus, binds to the TEAD protein Scalloped (Sd) and turns on transcriptional targets (Goulev et al., 2008; Wu et al., 2008; Zhang et al., 2008). The majority of these targets promote organ growth by suppressing apoptosis and stimulating cell growth and cell proliferation (Harvey and Tapon, 2007).

A key control step of the Hippo pathway is the localisation and kinase activity of Hippo. In epithelial cells, the scaffold protein Salvador promotes Hippo kinase activity by localising Hippo to the plasma membrane (Yin et al., 2013) and by inhibiting a large protein complex called the STRIPAK (Striatin-interacting phosphatase and kinase) complex (Bae et al., 2017). The STRIPAK complex contains PP2A as active phosphatase, which dephosphorylates a key Hippo auto-phosphorylation site and thus inhibits Hippo activity (Ribeiro et al., 2010; Zheng et al., 2017). dRassf can promote this recruitment of STRIPAK to Hippo and thus inactivate Hippo (Polesello et al., 2006; Ribeiro et al., 2010). Furthermore, the Hippo pathway can also be regulated downstream by membrane localisation of the kinase Warts by Merlin binding, which promotes Warts phosphorylation by Hippo and thus activation of the pathway (Yin et al., 2013). Finally, mechanical stretch of the epithelial cell cortex was shown to directly inhibit the Hippo pathway, likely mediated by the spectrin network at the cortex, promoting nuclear localisation of Yorkie (Fletcher et al., 2018; 2015). Despite this detailed knowledge about Hippo regulation in proliferating epithelial cells, little is known about how the Hippo pathway is regulated during post-mitotic muscle development and how it impacts muscle growth.

Here, we employ a systematic *in vivo* muscle-specific RNAi screen and identify various components of the Hippo pathway as essential post-mitotic regulators of flight muscle morphogenesis. We find that loss of Dlg5 or of the STRIPAK complex member Slmap, which interacts with Dlg5, as well as loss of the transcriptional regulator Yorkie results in too small muscles. These small muscles express lower levels of sarcomeric proteins and as a consequence contain fewer and defective myofibrils. Conversely, over-activation of Yorkie, either by removing the negative regulators Hippo or Warts or by enabling constitutive nuclear entry of Yorkie results in premature and excessive expression of sarcomeric proteins and consequently in chaotic myofibril assembly. Therefore, our findings suggest that the Hippo pathway contributes to the precise timing of sarcomeric gene expression and thus can generate a feedback mechanism for muscles to precisely coordinate sarcomeric protein levels during myofibril assembly and myofibril maturation. This provides an attractive mechanism for how regulated transcription can coordinate muscle growth with myofibril morphogenesis.

## Results

### Growth of *Drosophila* flight muscles

We chose the *Drosophila* indirect flight muscles to investigate post-mitotic muscle fiber growth. These muscles consist of two groups, the dorsal-longitudinal flight muscles (DLMs) and the dorso-ventral flight muscles (DVMs). Both groups form in the second thoracic segment during pupal development and despite differences during myoblast fusion and myotube attachment determining their location in the thorax, their development after 24 h after puparium formation (APF) is very similar (Dutta et al., 2004; Fernandes et al., 1991; Schönbauer et al., 2011). Thus, we focused our studies on the DLMs and for simplicity call them flight muscles in the remainder of the manuscript.

In order to quantify muscle fiber growth we measured fiber length and cross-sectional area of wild-type DLM flight muscles. At 24 h APF (at 27 °C) myoblast fusion is largely finished (Weitkunat et al., 2014), and the fibers have a length of about 270 μm and a cross-sectional area of about 1000 μm^2^ (Figure 1A,C, Supplementary Table 1). Then, flight muscles build up mechanical tension, compact to about 220 μm in length, while their diameter grows to about 2000 μm^2^, and assemble the immature myofibrils at 32 h APF (Lemke et al., 2019; Weitkunat et al., 2014) (Figure 1A,C). After 32 h, flight muscles grow about 2.5 times in length while keeping the same diameter until 48 h APF. After 48 h APF, they grow further to about 800 μm in length while increasing in diameter to almost 4000 μm^2^ until 90 h APF, which is shortly before eclosion at 27 °C (Figure 1A, Supplementary Table 1) (Spletter et al., 2018). Thus, in total, the volume of the individual muscle fibers increases more than 10-fold in less than 3 days (Figure 1A, Supplementary Table 1). Thus, indirect flight muscles are a good model to study rapid post-mitotic muscle fiber growth.

**Figure 1.**
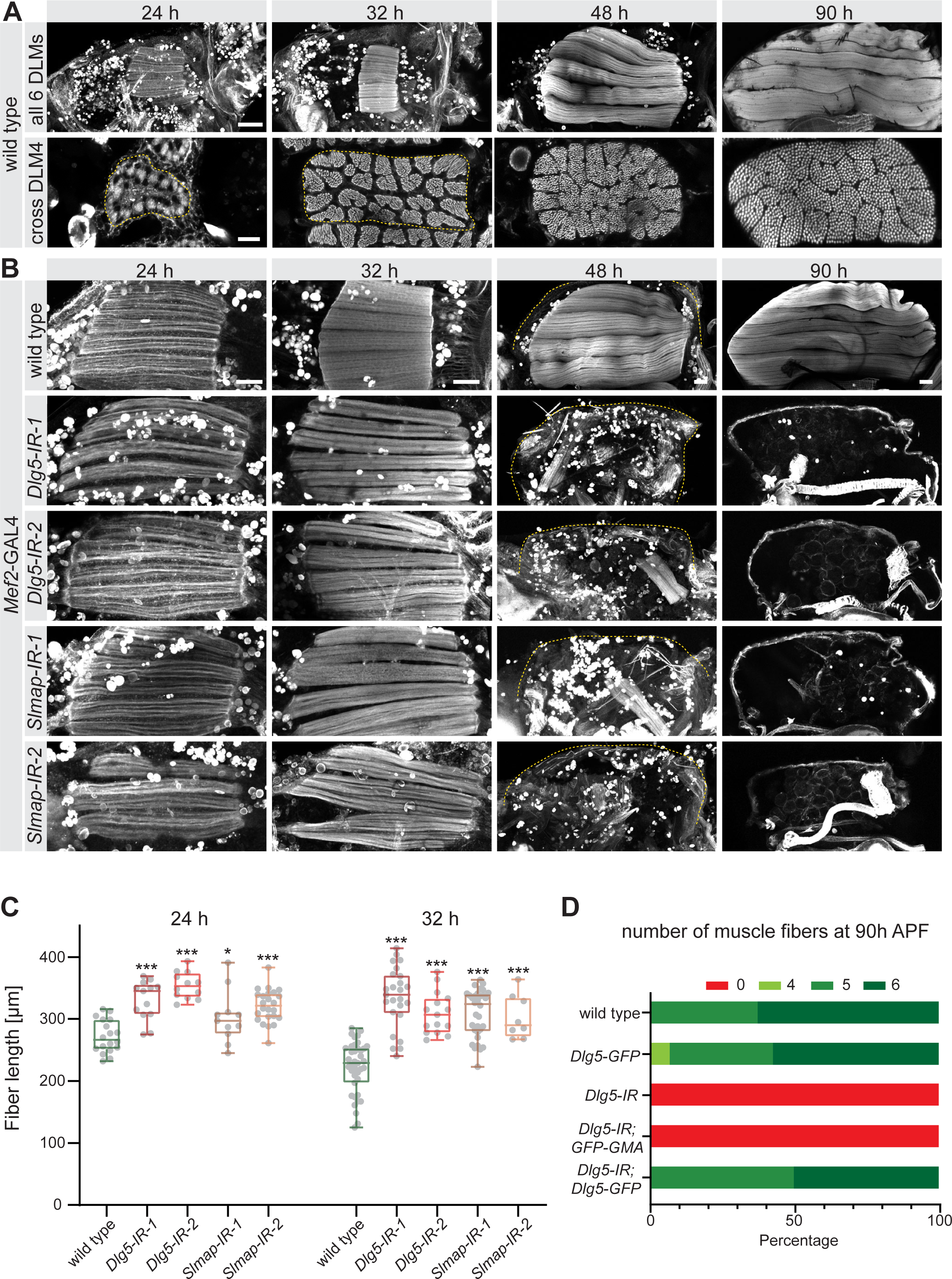
*Dlg5* and *Slmap* are essential for flight muscle morphogenesis. **A**. Time-course of wild-type dorsal longitudinal indirect flight muscle (DLM) development. Longitudinal sections (upper panel) of all DLMs and cryo cross-sections (lower panel) of dorsal longitudinal muscle 4 (DLM4) were stained for actin. Note the muscle fiber growth in length and width. Scale bars represent 100 μm for longitudinal and 10 μm for cross-sections. **B**. Longitudinal views of developing flight muscles at 24 h, 32 h, 48 h and 90 h APF of wild type, *Dlg5* or *Slmap* knockdown genotypes (independent RNAi lines *IR-1* and *2*) stained for actin. Note that *Dlg5-IR* and *Slmap-IR* muscles are too long at 24 h and 32 h APF, and are lost after 32 h APF. The dotted lines highlight the cuticle. Scale bars represent 50 μm. **C**. Box plot showing flight muscle fiber length at 24 h and 32 h APF of the indicated genotypes. Each dot represents the average muscle length from one pupa (see Methods). Box extends from 25% to 75%, line marks median, whiskers extend from max to min, all data points superimposed. Student’s t-test, *** p-value <0.001, *p value <0.05. All following box plots are plotted the same way. n ≥8 pupae for each genotype. **D**. Number of flight muscle fibers in half thoraces at 90 h APF of the indicated genotypes. Note that *UAS-Dlg5-GFP* but not *UAS-GFP-GMA* rescues the fiber atrophy phenotype of *Dlg5* knockdown (*Dlg1-IR-1*, in all the following figures *IR* refers to *IR-1*).

### *Dlg5* and *Slmap* are essential for flight muscle morphogenesis

To identify regulators of muscle growth we have investigated genes identified in a genome-wide muscle-specific RNAi study that had resulted in flightless or late developmental lethality when knocked-down using muscle-specific *Mef2*-GAL4 (Schnorrer et al., 2010). Our analysis identified two genes *Dlg5* (*Discs large 5, CG6509*) and *Slmap* (*Sarcolemma associated protein, CG17494*), which are conserved from *Drosophila* to human and when knocked-down using several independent RNAi constructs result in viable but completely flightless flies (Figure 1 supplement 1, Supplementary Table 1). Inspection of the thoraces of these animals revealed complete flight muscle atrophy in pupae at 90 h APF (Figure 1B). Expression of a UAS-*Dlg5-GFP* but not a UAS-GFP-Gma control construct, was able to rescue the number of muscle fibers of *Dlg5* knock-down (*Dlg5-IR-1*) flies to wild type providing further strong evidence for the specificity of the knock-down phenotype (Figure 1D). We conclude that *Dlg5* and *Slmap* are two conserved genes essential for flight muscle morphogenesis during pupal stages.

To identify the developmental time point when *Dlg5* and *Slmap* are required, we analysed pupal stages and found that at 24 h APF all flight muscles are present after *Dlg5* or *Slmap* knockdown. However, the fibers are more than 20% longer than wild type and fail to compact at 32 h APF when myofibrils normally assemble (Figure 1B,C, Supplementary Table 1). Interestingly, after 32 h APF, when wild-type myofibers strongly grow in length, *Dlg5* and *Slmap* knock-down fibers undergo complete flight muscle atrophy until 48 h APF (Figure 1B). Taken together, these data demonstrate that *Dlg5* and *Slmap* play an essential role during stages of myofibril assembly and muscle fiber growth.

### Dlg5 interacts with Slmap, a STRIPAK complex member, in *Drosophila* muscle

To define a molecular mechanism for how Dlg5 regulates muscle morphogenesis we performed GFP immunoprecipitation followed by mass-spectrometry, using the functional *UAS-Dlg5-GFP*, which was expressed in pupal muscles with *Mef2*-GAL4. Interestingly, we not only identified Slmap as a binding partner of Dlg5 in muscle, but also Fgop2, GckIII, Striatin (Cka), and the catalytic subunit of PP2A phosphatase (Mts) (Figure 2A,B, Supplementary Table 1). All these proteins are members of the STRIPAK complex and have been described to interact closely in mammals (Hwang and Pallas, 2014). This suggests that the composition of the STRIPAK complex in fly muscles is similar to mammals (Figure 2C), and Dlg5 is either a core member or closely interacts with this complex in muscle.

**Figure 2.**
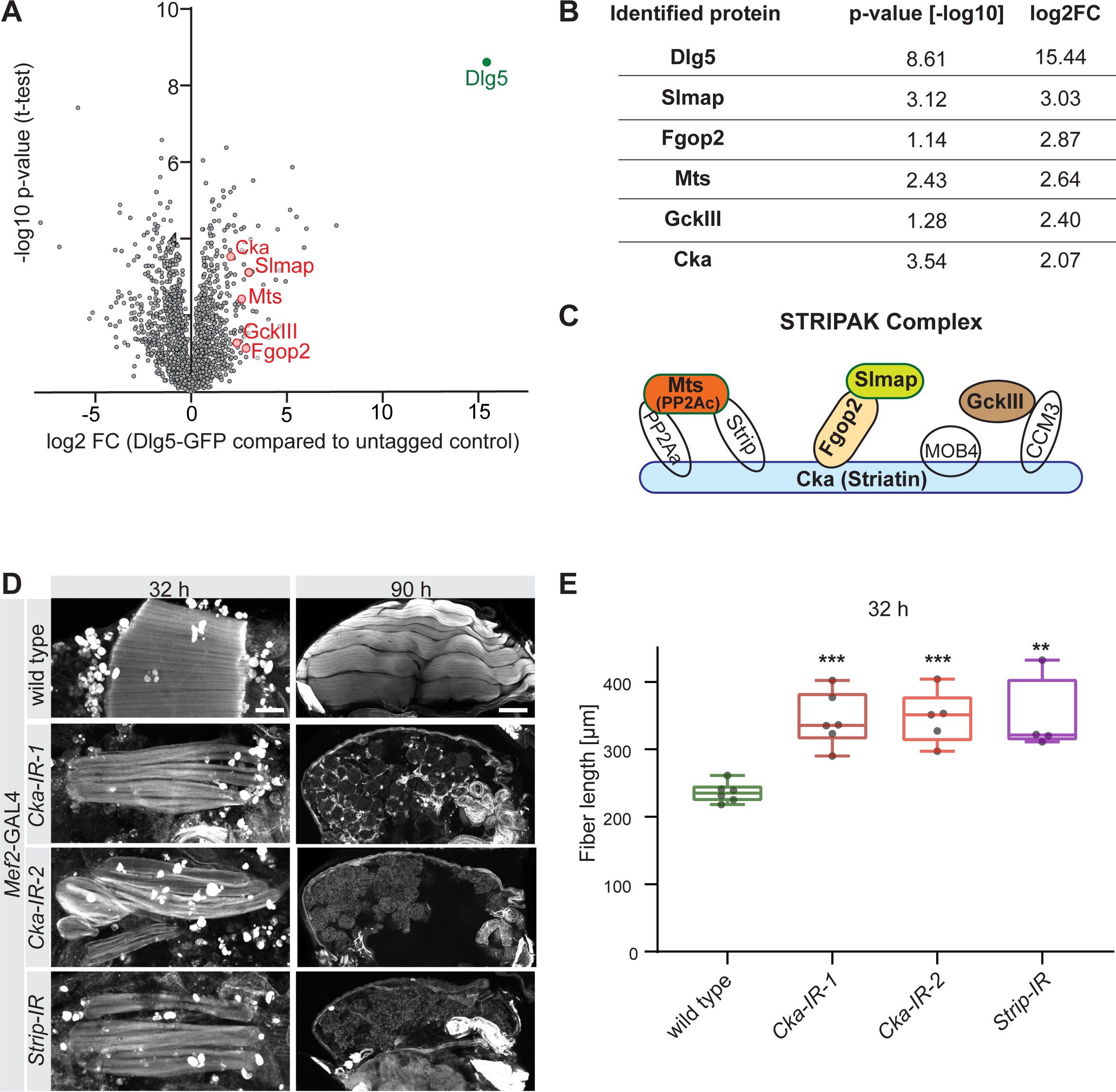
Dlg5 binds to the STRIPAK complex. **A**. Volcano plot showing proteins enriched in GFP pull-down from *Mef2*-GAL4, *UAS-Dlg5-GFP* 24 h - 48 h APF pupae. Y-axis shows statistical significance (-log10) and x-axis the log_2_ fold change (FC) compared to *Mef2*-GAL4 control (see Supplementary Table 1). STRIPAK complex components are highlighted in red. **B**. Table showing the log_2_ fold change and p-values of the selected STRIPAK complex components. **C**. Simplified schematic of the STRIPAK complex (adapted from (Duhart and Raftery, 2020)). Proteins identified in the Dlg5-GFP pull-down are coloured. **D**. Flight muscles at 32 h and 90 h APF of wild type, *Cka* (independent RNAi lines *IR-1* and *2*) or *Strip* knockdown genotypes stained for actin. Note the compaction defect at 32 h APF followed by muscle atrophy. Scale bars represent 50 μm for 32 h and 100 μm for 90 h APF. **E**. Box plot showing DLM fiber length at 32h APF of the indicated genotypes. Each dot is the average from one pupa. Student’s t-test, *** p-value <0.001, ** p-value <0.01.

To functionally test if members of the STRIPAK complex other than Dlg5 and Slmap play a similarly important role in flight muscles, we knocked-down various component members and found that knock-down of *Striatin* (*Cka*) and *Strip* indeed result in very similar phenotypes to *Dlg5* and *Slmap* knock-down: flight muscles at 32 h APF are longer than wild type, suggesting a fiber compaction defect, and undergo muscle atrophy after 32 h APF (Figure 2D,E). Together, these data show that Dlg5 interacts with the STRIPAK complex in flight muscles, of which several members including Slmap are important for flight muscle morphogenesis.

### The Hippo pathway regulates the developmental timing of muscle morphogenesis

As the STRIPAK complex was shown to dephosphorylate and thus inactivate Hippo (Ribeiro et al., 2010; Zheng et al., 2017), we wondered if the muscle morphogenesis phenotype we observed could be linked to a function of the Hippo pathway in growing flight muscles. When knocking-down the Hippo pathway transcriptional co-activator *yorkie* in muscles, we observed a failure of muscle compaction at 32 h APF, flight muscle atrophy at 48 h APF and consequently flightless adults (Figure 3A,B, Figure 3 supplement 1A, Supplementary Table 1). The myofiber compaction defect of *yorkie* knock-down muscles is corroborated by fiber cross-sections at 24 h and 32 h APF revealing much thinner muscles and phenocopying the *Dlg5* and *Slmap* loss of function (Figure 3 supplement 1B).

**Figure 3.**
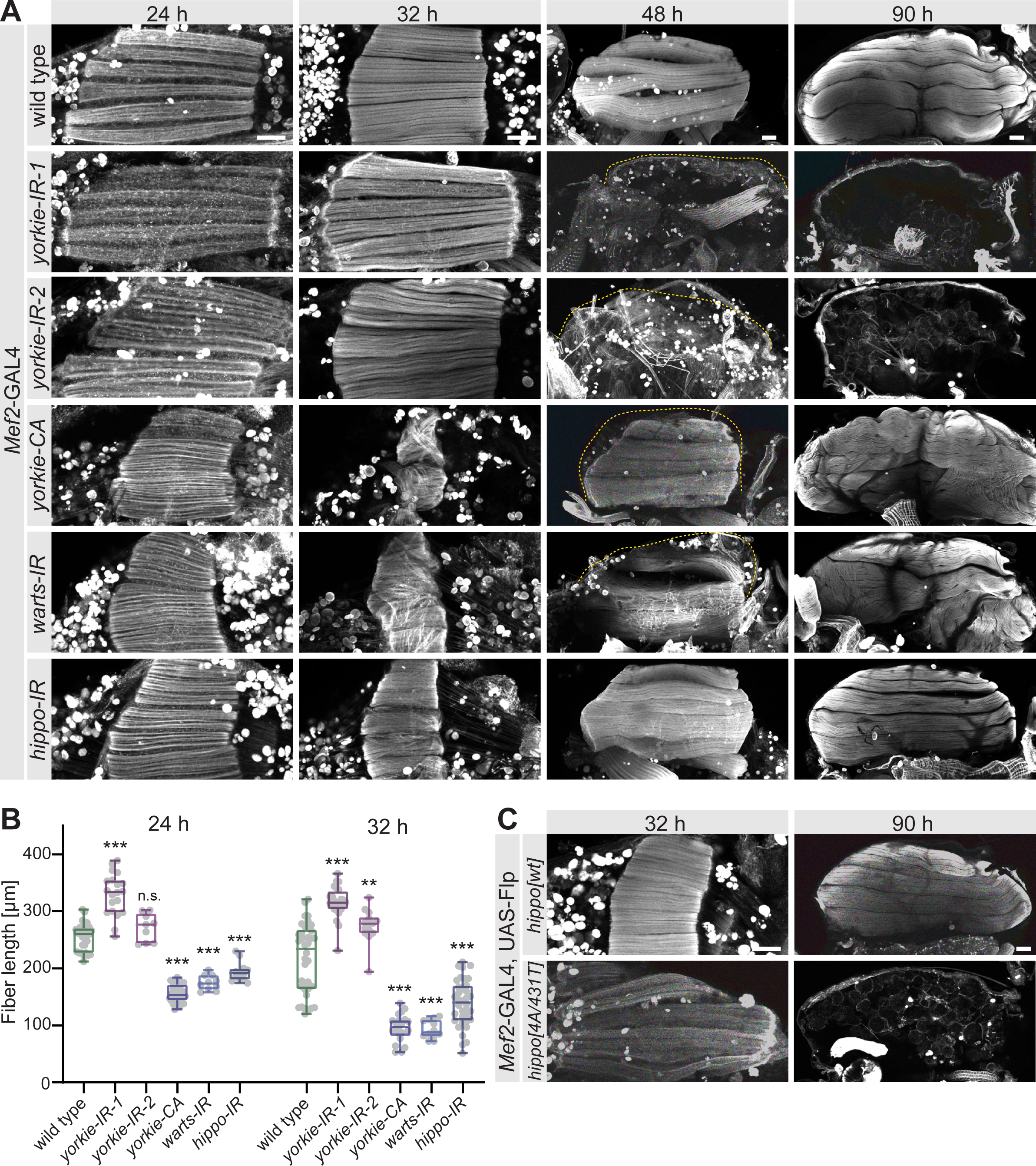
The Hippo pathway regulates muscle morphogenesis. A. Flight muscles at 24 h, 32 h, 48 h and 90 h APF from wild type, *yorkie* knock-down (independent RNAi lines *IR-1* and *2*), *yorkie-CA*, as well as *warts* and *hippo* knockdown genotypes stained for actin. The dotted lines highlight the cuticle. Note the too long *yorkie-IR* muscles but too short *yorkie-CA, warts-IR* and *hippo-IR* muscles at 24 h and 32 h APF. **B**. Box plot showing muscle fiber length at 24 h and 32 h APF of the indicated genotypes. Student’s t-test, *** p-value<0.001, ** p value<0.01. **C**. Flight muscles at 32 h and 90 h APF from pupae expressing either wild-type *hippo* or Slmap-binding deficient *hippo[4A/ 431T]* (under control of the milder tubulin promoter). The FRT stop cassette is removed by muscle specific *Mef2*-GAL4 driven Flp recombinase. All scale bars represent 50 μm.

In contrast to the myofiber compaction defect of *yorkie* knock-down muscles, expression of an activated form of Yorkie (*yorkie-CA*), which cannot be phosphorylated by Warts, results in premature muscle fiber compaction already at 24 h APF and strongly hyper-compacted muscle fibers at 32 h APF (Figure 3A,B). The increased cross-sectional area is particularly obvious in cross-sections of *yorkie-CA* fibers (Figure 3 supplement 1B). Importantly, we observed the same phenotypes after knock-down of each of the two kinases *hippo* and *warts*, both negative regulators of Yorkie nuclear entry (Figure 3A,B). This strongly suggests that the Hippo pathway, by regulating phosphorylation of the transcriptional co-activator Yorkie, is essential for the correct developmental timing of flight muscle morphogenesis: too much active Yorkie accelerates myofiber compaction, while too little active Yorkie blocks it.

To further corroborate that the STRIPAK complex regulates the Hippo pathway in flight muscles we used a recently characterised Hippo construct (*hippo[4A/431T]*), which lacks the four auto-phosphorylation sites in Hippo required to bind to the STRIPAK phosphatase complex via Slmap. This Hippo[4A/431T] protein cannot be dephosphorylated on regulatory T195 by STRIPAK and thus is constitutively active (Zheng et al., 2017). Interestingly, expression of *hippo[4A/431T]* in muscle after *Mef2*-GAL4 driven flip-out also resulted in a muscle fiber compaction defect at 32 h APF and muscle atrophy thereafter, phenocopying the STRIPAK and *yorkie* loss of function phenotypes (Figure 3C). Taken all these data together, we conclude that Dlg5 and members of the STRIPAK complex are key regulators of the Hippo pathway, which controls the developmental timing of flight muscle morphogenesis in *Drosophila*.

### The Hippo pathway is required post-mitotically in flight muscle fibers

Indirect flight muscles are formed by fusion of several hundred myoblasts until 24 h APF (Weitkunat et al., 2014). These myoblasts emerge during embryonic development and proliferate extensively during larval stages (Bate et al., 1991; Gunage et al., 2014; Roy and VijayRaghavan, 1998). As knock-down of *Dlg5*, STRIPAK complex and Hippo pathway members with *Mef2*-GAL4 results in flightlessness and not in lethality (except for *warts*, see Figure 3 - supplement 1), it is unlikely that general myoblast proliferation during larval stages is affected, which would result in defects of all adult muscles. However, since *Mef2*-GAL4 is already active during larval stages, we wanted to exclude that the observed muscle phenotypes are caused by myoblasts proliferation defects during larval stages. Hence, we conditionally activated GAL4 only during pupal stages using temperature sensitive GAL80 (GAL80ts, see Methods)(McGuire et al., 2003) and quantified myoblast fusion rates by counting the nuclei of dorsal longitudinal flight muscle 4 (DLM4). We found comparable numbers of nuclei at 24 h APF ruling out a major contribution of myoblast proliferation or fusion to the phenotype (Figure 4 supplement 1).

Importantly, these GAL80ts *Mef2*-GAL4 *Dlg5* and *yorkie* knock-down muscles do display the same fiber compaction defect as observed with *Mef2*-GAL4 resulting in longer but thinner fibers with grossly comparable volumes at 24 h APF (*yorkie-IR* is slightly smaller) (Figure 4A,B). These fibers do not compact at 32 h APF and undergo atrophy leading to no remaining fibers at 48 h or 90 h APF (Figure 4C, D), phenocopying the constitutive knock-down of *Dlg5* or *yorkie*. Conversely, conditional expression of yorkie-CA during pupal stages results in premature compaction at 24 h APF and very short fibers at 32 h APF that grow to disorganised fibers at 90 h APF (Figure 4A-D). These phenotypes resemble the constitutive *Mef2*-GAL4 driven phenotypes demonstrating a role for *Dlg5* and *yorkie* in muscle fibers during pupal stages.

**Figure 4.**
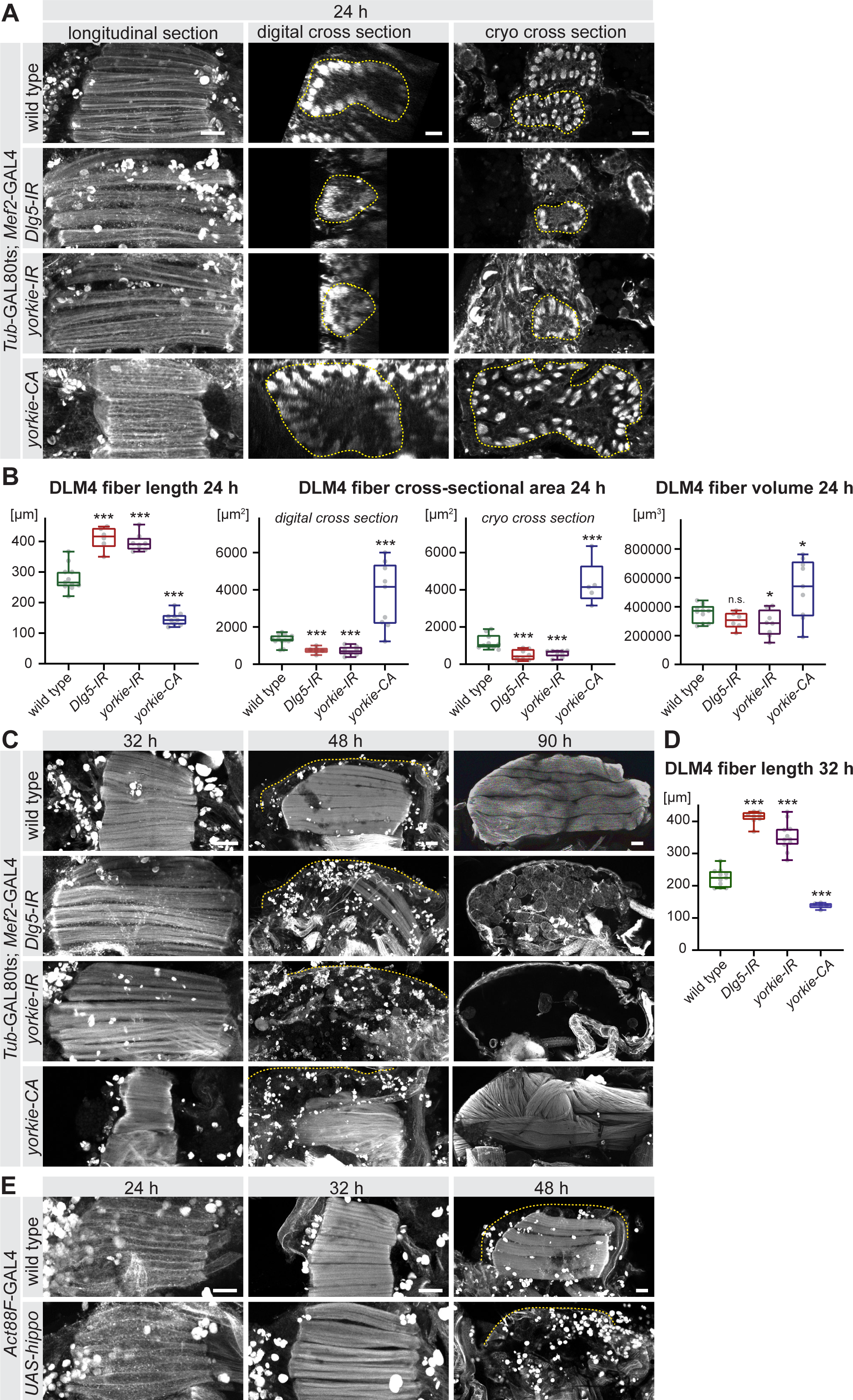
The Hippo pathway is important for post-mitotic muscle development. **A-D**. Developing flight muscles at 24 h, 32 h, 48 h and 90 h APF from wild type, *Dlg5-IR, yorkie-IR* and *yorkie-CA Tub*GAL80ts *Mef2*-GAL4 pupae, in which GAL4 activity was restricted to pupal stages (shift to 31°C at 0 h APF). **A**. Longitudinal sections of all flight muscles, as well as digital and cryo cross-sections of dorsal longitudinal muscle 4 (DLM4) at 24 h APF. Dotted lines highlight the fiber area. **B**. Box plot displaying DLM4 muscle fiber length, fiber cross-sectional area from digital and cryo sections as well as the calculated DLM4 fiber volume at 24h APF (calculated by multiplying length with cross-sectional area from digital cross-sections for each pupa). Student’s t-test, *** p-value<0.001, * p-value<0.05. **C**. Flight muscles at 32 h, 48 h and 90 h APF. Note the too long muscles in *Dlg5-IR, yorkie-IR* at 32 h APF followed by muscle atrophy and the too short *yorkie-CA* fibers at 32 h that develop disorganised fibers at 90 h APF. **D**. Box plot illustrating the average muscle fiber length at 32 h APF. Student’s t-test, *** p-value<0.001. **E**. Control and *Act88F*-GAL4 *UAS-hippo* flight muscles at 32 h and 90 h APF. Note the induced fiber compaction defect followed by the muscle atrophy. All scale bars represent 50 μm for longitudinal sections and 10 μm for cross-sections.

To further corroborate a post-mitotic role of the Hippo pathway in muscle fibers, we over-expressed *hippo* with the strong, strictly post-mitotic flight muscle specific driver *Act88F*-GAL4, which is only active after myoblast fusion (Bryantsev et al., 2012; Spletter et al., 2018). This post-mitotic over-expression of *hippo* resulted in flight muscle compaction defects at 32 h APF and muscle atrophy at 48 h APF (Figure 4E). Together, these data demonstrate that the Hippo pathway and its regulator Dlg5 are required post-mitotically in flight muscle fibers for the correct timing of morphogenesis.

### The Hippo pathway regulates post-mitotic muscle fiber growth

The wild-type flight muscle fibers grow in volume from 24 h to 48 APF (see Figure 1), while *yorkie* or *Dlg5* knock-down fibers undergo atrophy after 32 h APF. As the Yorkie activity is known to suppress apoptosis in epithelial tissues (Harvey and Tapon, 2007) we asked if we could rescue fiber atrophy by over-expressing the apoptosis inhibitor Diap1. Indeed, over-expression of Diap1 during pupal stages in GAL80ts *Mef2*-GAL4 (hereafter abbreviated as GAL80ts) *yorkie* and *Dlg5* knock-down fibers substantially rescues fiber atrophy, often resulting in the normal number of six muscle fibers at 48 h APF (Figure 5A, compare to Figure 4C). This demonstrates that apoptosis contributes to flight muscle fiber atrophy in *yorkie* and *Dlg5* knock-down muscles.

**Figure 5.**
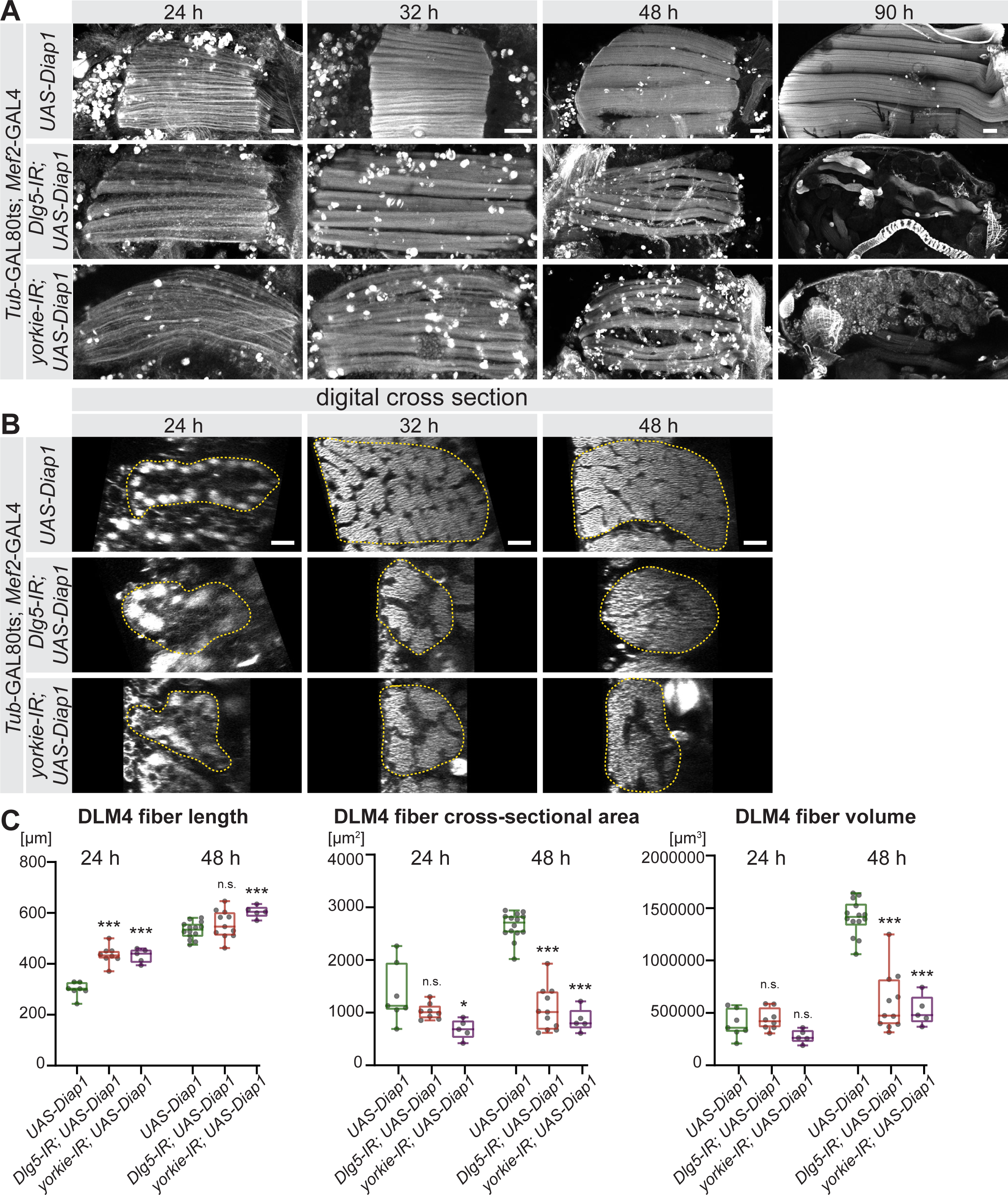
The Hippo pathway is essential for post-mitotic muscle fiber growth. **A, B**. Longitudinal sections of all DLMs (A) and digital cross-sections of DLM4 (B) at 24 h, 32 h, 48 h and 90 h APF from wild type, *Dlg5-IR, yorkie-IR* and *yorkie-CA Tub*GAL80ts *Mef2*-GAL4 (shifted to 31°C at 0 h APF) stained for actin. Scale bars represent 50 μm for longitudinal sections and 10 μm for cross-sections. In B dotted lines highlight the fiber area. **C**. Box plots showing DLM4 fiber length, digital cross-sectional area and volume (calculated by multiplying length with cross-sectional area for each pupa) at 24 h and 48 h APF. Student’s t test, *** p-value <0.001, * p-value <0.05.

Presence of muscle fibers at 48 h APF enabled us to quantitatively investigate the role of the Hippo pathway during the post-mitotic muscle fiber growth. As in Figure 4, we used digital cross-sections of large confocal stacks to quantify the cross-sectional area of dorsal longitudinal flight muscle 4 (DLM4) and together with the fiber length calculated the fiber volume (Figure 5). Similar to what we showed in Figure 4, GAL80ts *Diap1* expressing control fibers have a comparable volume to GAL80ts *Diap1* expressing *yorkie* and *Dlg5* knock-down fibers at 24 h APF (Figure 5A-C), showing that they start into the muscle growth phase with comparable sizes.

However, until 32 h APF, *Diap1* control muscles increase their cross-sectional area followed by growth in length until 48 h APF to increase their volume about 4-fold within 24 h (Figure 5A-C). Interestingly, GAL80ts *Diap1* expressing *yorkie* or *Dlg5* knock-down muscles fail to normally increase their cross-sectional area at 32 h and 48 h APF, thus resulting in smaller volumes at 48 h APF (Figure 5A-C). These thin muscles do not survive until 90 h APF despite over-expression of the apoptosis inhibitor Diap1 (Figure 5A). Taken together, these data provide strong evidence that the Hippo pathway and its transcriptional co-activator Yorkie are required to enable normal post-mitotic growth of flight muscle fibers, likely by regulating the developmental timing of muscle morphogenesis.

### The Hippo pathway is essential for myofibrillogenesis

To investigate the molecular mechanism of the muscle growth defect in detail, we quantified myofibrillogenesis in these muscles. Control *Diap1* expressing muscles have assembled immature myofibrils at 32 h APF (Weitkunat et al., 2014). These immature myofibrils are continuous and thus can be easily traced throughout the entire field of view (Figure 6A,B, Figure 6 supplement 1A). In contrast, GAL80ts *Diap1* expressing *yorkie* and *Dlg5* knock-down muscles fail to properly assemble their myofibrils at 32 h APF resulting in only short myofibril traces (Figure 6A,B, Figure 6 supplement 1A). Concomitant with the myofibril assembly defect, we also found that the spacing of the nuclei is defective. In control muscle fibers the nuclei are present mainly as single rows located between myofibril bundles, whereas in *yorkie* and *Dlg5* knock-down muscles they form large centrally located clusters (Figure 6 supplement 1B). This indicates that at 32 h APF, Hippo signalling is required within the muscle fibers to trigger proper myofibril assembly and nuclear positioning.

**Figure 6.**
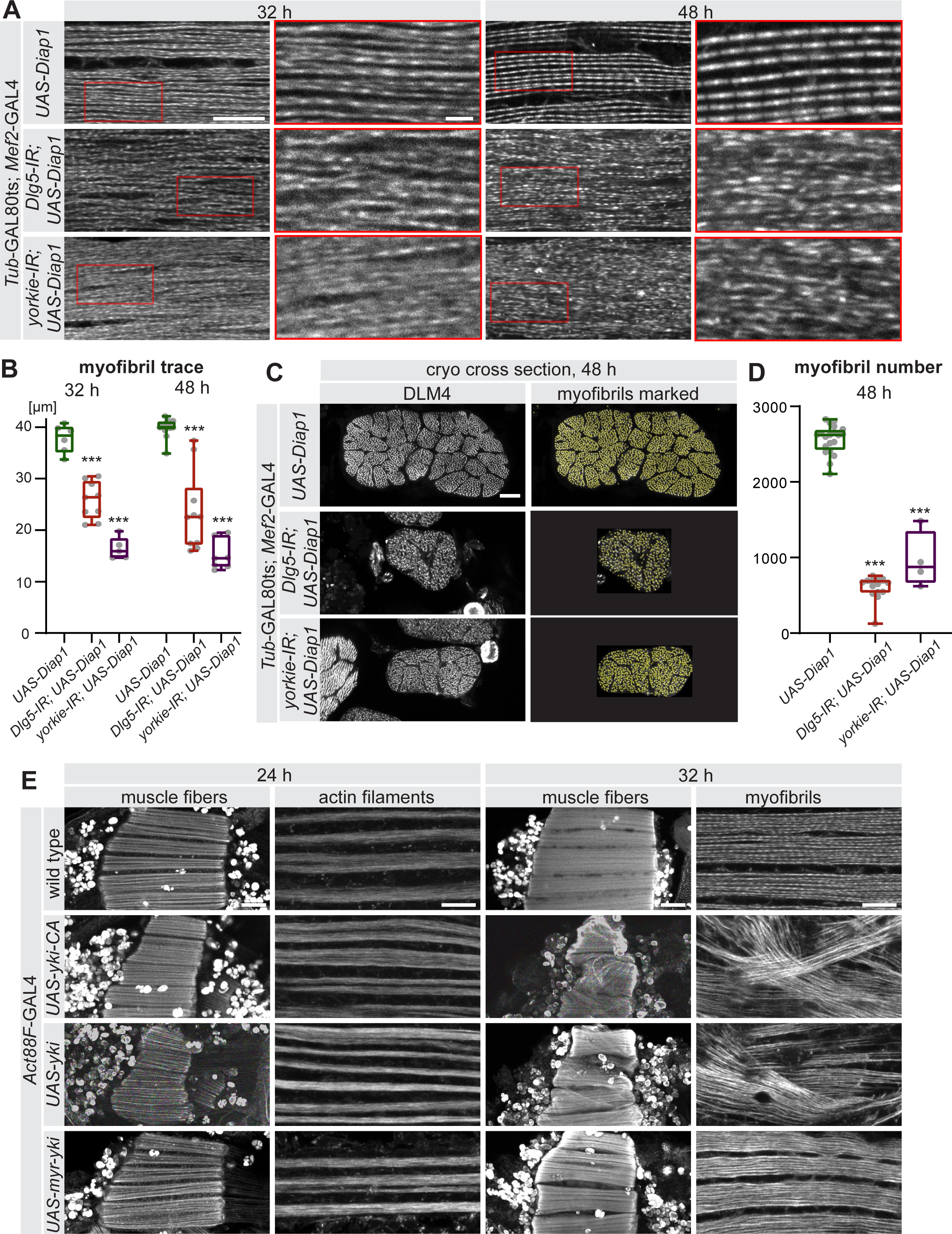
The Hippo pathway is essential for myofibrillogenesis. **A**. Myofibrils visualised by phalloidin from control, *Dlg5-IR* and *yorkie-IR UAS-Diap1 Tub*GAL80ts *Mef2*-GAL4 muscles at 32 h and 48 h APF (shifted to 31°C at 0 h APF). The red-boxed areas are magnified. Note the less regular myofibril pattern of *Dlg5-IR* and *yorkie-IR* muscles at 32 h APF that makes it hard to trace an individual myofibril (see Figure 6 supplement 1A). Even at 48 h APF, myofibrils from *Dlg5-IR* and *yorkie-IR* are hard to trace continuously. Scale bars represent 10 μm in the overviews and 2 μm in the zoomed red boxes. **B**. Box plot of traced myofibril length in a 40 × 20 × 2.5 μm volume (see Figure 6 supplement 1A). Student’s t test, *** p-value <0.001. **C**. Cryo cross-sections of DLM4 from control, *Dlg5-IR* and *yorkie-IR UAS-Diap1 Tub*GAL80ts *Mef2*-GAL4 muscles at 48 h APF (shifted to 31°C at 0 h APF). Yellow dots represent the myofibrils recognized by the MyofibrilJ plug-in to automatically count the number of myofibrils per DLM4 fiber (Spletter et al., 2018). Scale bar represents 10 μm. **D**. Box plot of myofibril number in DLM4 of indicated genotypes at 48 h APF. Student’s t test, *** p-value <0.001. **E**. Flight muscle and myofibril morphologies of wild-type muscles expressing *yorkie-CA, yorkie* or *myr-yorkie* under the control of post-mitotic *Act88F*-GAL4 at 24 h and 32 h APF. Scale bars represent 50 μm in muscle fiber images and 10 μm in myofibril images.

The myofibril defect becomes even more pronounced at 48 h APF when control myofibrils have matured and sarcomeres are easily discernable (Figure 6A), while no organised sarcomeres are present in GAL80ts *yorkie* and *Dlg5* knock-down muscles and myofibril traces remain short (Figure 6A,B, Figure 6 supplement 1A). Furthermore, cryo cross-sections revealed that not only the cross-sectional area but also the total number of myofibrils is strongly reduced in GAL80ts *yorkie* and *Dlg5* knock-down muscles compared to control (Figure 6C,D, Figure 6 supplement 1C). These data demonstrate that the Hippo pathway controls both the morphological quality of the myofibrils at the assembly and maturation stages as well as their quantity. As myofibrils occupy most of the muscle fiber space, their reduced amount likely causes the reduced muscle size in *Dlg5* or *yorkie* knock-down fibers.

### Yorkie is a transcriptional co-regulator in muscle fibers

It was recently shown that the transcription of most sarcomere key components is tightly regulated starting shortly before myofibril assembly and being strongly boosted during myofibril maturation (Spletter et al., 2018). Thus, we reasoned that Yorkie activity may be involved in this transcriptional regulation step to control the timing of myofibrillogenesis. However, it had also been recently shown that Yorkie can regulate myosin contractility directly at the cell membrane without entering into the nucleus (Xu et al., 2018). As we have thus far failed to unambiguously locate Yorkie protein in muscle fibers during development, we used genetic tools to address this important point. To test whether Yorkie may play a role outside of the nucleus, we manipulated Yorkie levels by over-expressing different Yorkie variants post-mitotically using *Act88F*-GAL4 and investigated the consequences at 24 h and 32 h APF. Over-expression of either Yorkie-CA, whose import into the nucleus is uncoupled from the Hippo pathway, or wild-type Yorkie, whose nuclear import is regulated by Hippo, both result in premature muscle fiber compaction at 24 h APF, with seemingly normal actin filaments (Figure 6E). Strikingly, the muscle fiber hyper-compaction at 32 h APF coincides with a chaotic organisation of the myofibrils, with many myofibrils not running in parallel but in various directions (Figure 6E, Figure 6 supplement 1D). This suggests that the hyper-compaction phenotype upon Yorkie over-expression is likely caused by uncontrolled and premature force production of the chaotically assembling myofibrils.

In contrast, over-expression of a membrane-anchored myristoylated form of Yorkie, which has been shown to activate myosin contractility at the epithelial cell cortex without going into the nucleus (Xu et al., 2018), does not result in premature muscle fiber compaction at 24 h APF. Furthermore, these muscles display normally oriented parallel myofibrils at 32h APF (Figure 6E). These results indicate that the observed myofibril and fiber compaction defects are caused by a transcriptional response of Yorkie in the nucleus.

This interpretation is corroborated by loss of function data of the transcriptional activator *scalloped* (*sd*), which is the essential transcriptional co-factor of Yorkie in the nucleus. Knock-down of *scalloped* results in severe muscle atrophy and no remaining muscles at 90 h APF (Figure 6 supplement 1E). Together, these genetic data suggest that Hippo signalling regulates Yorkie phosphorylation and thus its nuclear entry to trigger a transcriptional response that controls myofibril development and muscle fiber growth.

### The Hippo pathway controls expression of key sarcomere components

To investigate the transcriptional role of the Hippo pathway during flight muscle development we performed muscle-specific transcriptomics of wild-type flight muscles compared to different *yorkie* ‘loss of function’ (*Dlg5-IR, Slmap-IR* and *yorkie-IR*) and *yorkie* ‘gain of function’ (*yorkie-CA and hippo-IR*) conditions. We dissected flight muscles from 24 h and 32 h APF, isolated RNA and applied a sensitive 3-prime end mRNA sequencing method (BRB-seq)(Alpern et al., 2019), which handles small amounts of mRNA (see Methods). We found a clustering of biological replicates and similar genotypes using principle components analysis and comparable read count distributions across all samples (Figure 7 supplement 1). This verifies BRB-seq as a reliable method to quantitatively compare gene expression from small amounts of developing muscle tissue across multiple samples.

We applied the selection criteria log2FC > 1 and adjusted p-value < 0.05 to identify differentially expressed genes compared to wild type (Supplementary Table 2). Applying FlyEnrichr (Kuleshov et al., 2016) on the differential data sets, we found a strong enrichment for muscle and, in particular, for sarcomere and myofibril Gene Ontology terms (GO-terms) in the differentially expressed genes of all three *yorkie* ‘loss of function’ muscle genotypes (*Dlg5-IR, Slmap-IR* and *yorkie-IR*) at 24 h APF (Figure 7A, Supplementary Table 3). Importantly, expression of many core sarcomeric components, including both titin homologs *sallimus* (*sls*) and *bent* (*bt*), *Myosin heavy chain* (*Mhc*), *Myofilin* (*Mf*), *Paramyosin* (*Prm*), tropomyosins (*Tm1, Tm2*), flight muscle specific actin (*Act88F*) and *Obscurin* (*Unc-89*), as well as sarcomere dynamics regulators, including myosin phosphatase (*Mbs*), a flight muscle formin (*Fhos*) and the spektraplakin *shortstop* (*shot*) are consistently reduced in *yorkie* ‘loss of function’ muscle genotypes at 24 h APF (Figure 7B). Furthermore, expression of mRNAs coding for proteins linking the nuclei to the cytoskeleton, such as the Nesprin family members *klar* and *Msp300*, are also strongly reduced (Figure 7B), which may explain the observed nuclei position defect in *yorkie* and *Dlg5* knock-down muscles (Figure 6 supplement 1B). *Msp300* and *Prm* are amongst the only 6 genes that are significantly down-regulated in all three loss of function conditions at 24 h APF (Supplementary Table 2). This strongly suggests that nuclear entry of Yorkie contributes to the transcriptional induction of sarcomeric protein coding genes as well as genes important to link the nuclei to the sarcomeres. This transcriptional induction was shown to precede sarcomere assembly (Spletter et al., 2018) and thus may provide a molecular explanation of the observed flight muscle compaction and myofibril assembly defects of *Dlg5-IR, Slmap-IR* and *yorkie-IR* muscles.

**Figure 7.**
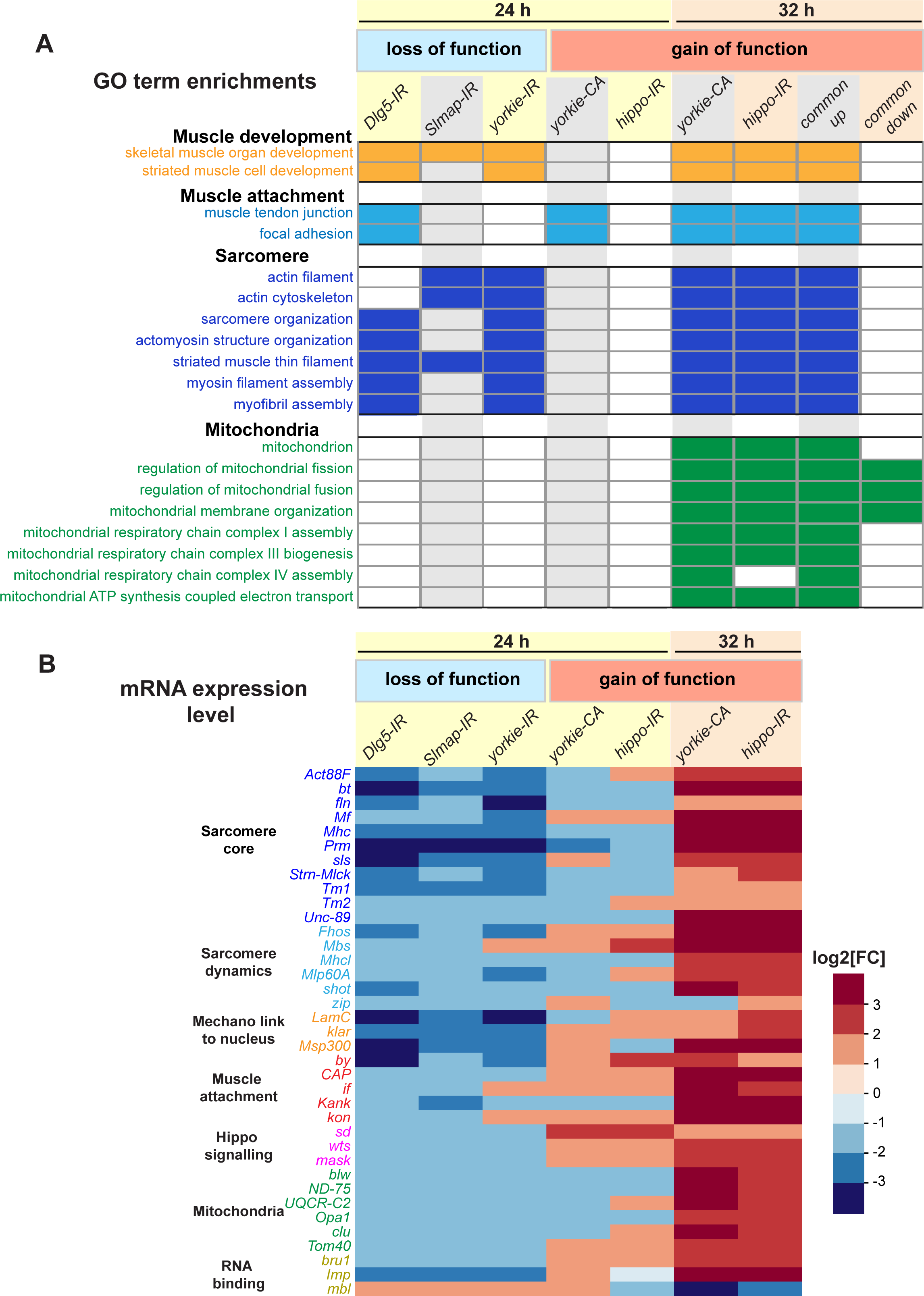
Yorkie transcriptionally controls sarcomeric and mitochondrial genes expression. **A**. Gene ontology (GO)-term enrichments in genes lists that are significantly changed in the various loss (*Dlg5-IR, Slmap-IR, yorkie-IR*) and gain of function (*yorkie-CA, hippo-IR*) yorkie conditions at 24 h and 32 h APF compared to wild-type controls (see Supplementary Table 3). Note the strong enrichment of sarcomere related GO-terms in the 24 h APF loss of function and the 32 h APF gain of function condition. 32 h APF gain of function is also strongly enriched for mitochondrial GO-terms. **B**. Plot displaying the log2-fold change of transcript levels for individual genes of the above genotypes compared to control (up in red, down in blue). Note the strong down-regulation of the sarcomeric genes in the 24 h APF loss of function conditions, in particular the titin homologs *bt* and *sls*, as well as *Mhc, Prm* and *Act88F*.

To complement the yorkie loss of function conditions, we also performed BRB-seq transcriptomics comparing *yorkie* ‘gain of function’ conditions (*yorkie-CA and hippo-IR*) to control. While we found few significantly differentially expressed genes at 24 h APF (including an induction of the transcriptional co-regulator *scalloped*) (Supplementary Table 2), we identified many significant changes in mRNA expression in *yorkie-CA* and *hippo-IR* myofibers at 32 h APF (Supplementary Table 2). Strikingly, sarcomeric core components and their regulators are amongst the top up-regulated genes on both lists (Figure 7B, Supplementary Table 2). Consequently, GO-term analysis of the differentially expressed genes identified a strong enrichment for sarcomere and myofibril GO-terms (Figure 7A, Supplementary Table 3). In addition to sarcomeric genes, genes important for myofibril attachment at muscle-tendon junctions, including the integrin attachment complex members *kon, if, CAP, by* and *Kank* are up-regulated in both gain of function genotypes (Figure 7B). Furthermore, we found an up-regulation of Hippo signalling regulators *mask*, which is required for efficient nuclear import of Yki (Sidor et al., 2019), and of the negative regulator *wts* (Figure 7B). This demonstrates that regulated Hippo activity is required to control Yorkie in order to tune expression of mRNAs coding for sarcomeric and myofibril attachment proteins.

During the stage of myofibril assembly, the mitochondria morphology changes and the expression of mitochondrial genes increase (Avellaneda et al., 2020; Spletter et al., 2018). Consistently, we found an induction of mitochondria dynamics and protein import regulators (Opa1, Tom40) in *hippo-IR and yorkie-CA* myofibers at 32 h APF as well as a consistent up-regulation of mRNAs coding for respiratory chain components, including the F1F0 ATP synthase complex (complex V) subunit *blw*, the NADH dehydrogenase (ubiquinone) subunit *ND-75* and the Ubiquinol-cytochrome c reductase subunit *UQCR-C2*, which are all required to boost ATP production during muscle fiber growth (Figure 7A,B, Supplementary Table 2). Taken together, these data strongly suggest that the Hippo pathway regulates the correct expression dynamics of many key muscle components, most prominently mRNAs coding for core sarcomeric and mitochondrial proteins to enable myofibril assembly and mitochondrial maturation.

### Yorkie controls sarcomeric protein dynamics

The most prominent phenotypes of the *yorkie* ‘loss of function’ group (*Dlg5-IR, Slmap-IR* and *yorkie-IR*) are defective muscle fiber compaction and severe myofibril assembly defects at 32 h APF. As the core sarcomeric proteins actin and myosin are required to assemble myofibrils and build up mechanical tension (Loison et al., 2018; Weitkunat et al., 2014) we chose to quantify protein levels of the major actin isoform in flight muscles Actin88F (Act88F) as well as the only *Drosophila* muscle Myosin heavy chain (Mhc). For both, we used GFP fusion proteins expressed under endogenous control (Sarov et al., 2016) and thus avoiding the variations often seen in antibody stainings (see Methods). Consistent with the transcriptomics data, we found a mild reduction of Mhc and Act88F protein levels in *yorkie* knock-down muscles at 24 h APF, which became more pronounced at 32 h APF (Figure 8A-C). This is consistent with the myofibril assembly defects found in *yorkie* knock-down muscles at 32 h APF.

**Figure 8.**
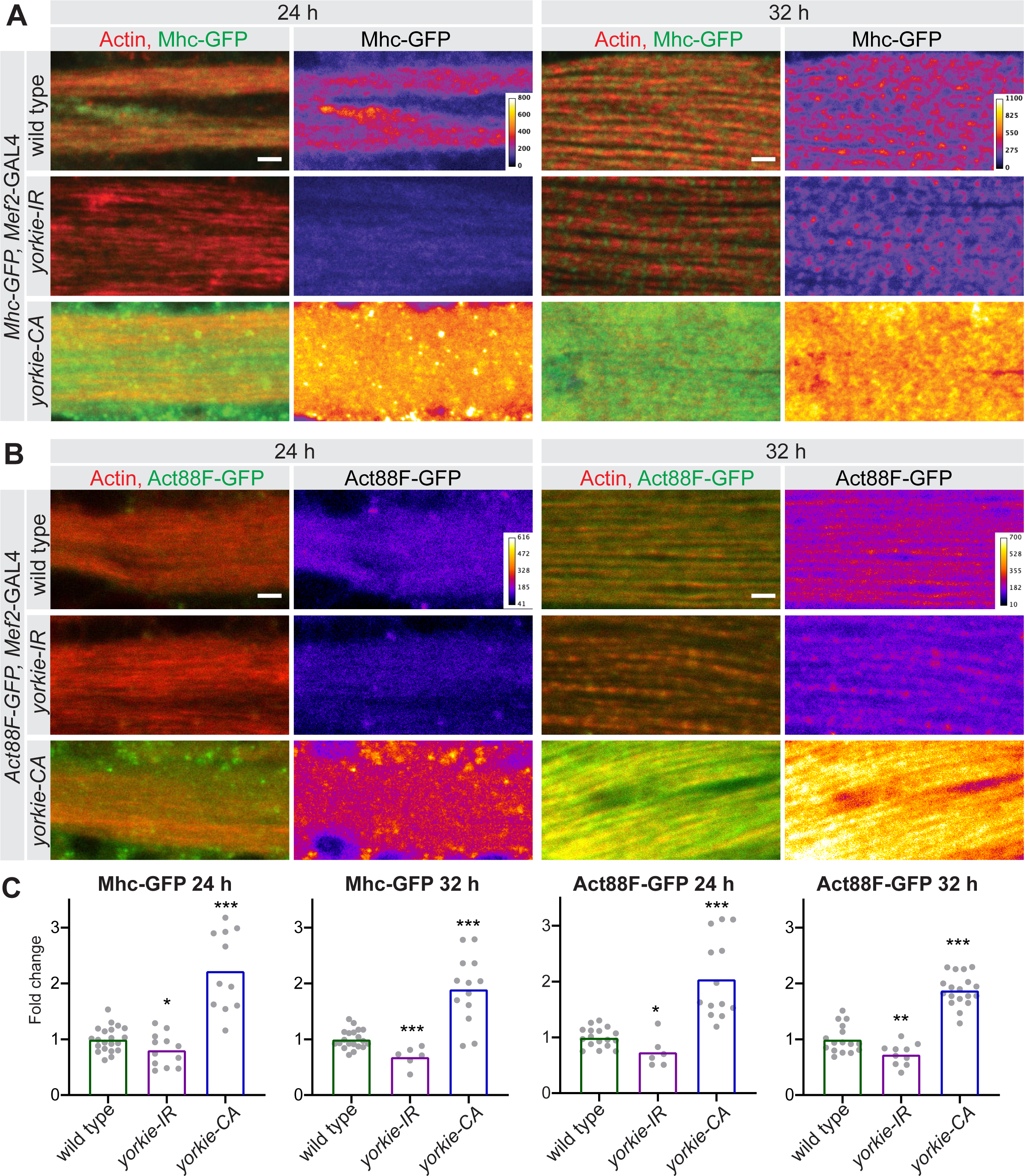
Yorkie regulates sarcomere protein expression. **A, B**. Mhc-GFP (A) and Act88F-GFP (B) proteins levels visualised with a GFP nanobody together with actin comparing wild-type control with *yorkie-IR* and *yorkie-CA* at 24 h and 32 h APF. Scale bars represent 2 μm. GFP channel is displayed using the “fire” look-up table. **C**. Quantification of relative GFP protein levels compared to wild type (wild-type mean set to 1 for each individual experiment). Bar plots display average normalised value and each dot represents one hemithorax. Student’s t test, *** p-value <0.001, ** p-value <0.01 * p-value <0.05.

Surprisingly, constitutive activation of *yorkie* (*yorkie-CA*) results in a boost of Mhc-GFP and Act88F-GFP protein expression already at 24 h APF, which is maintained at 32 h APF (Figure 8A-C). This increased acto-myosin expression may provide the molecular explanation of the premature compaction phenotype seen in *yorkie-CA* muscles at 24 h APF. Despite being expressed at high levels already at 24 h APF, both proteins fail to prematurely assemble into periodic myofibrils in *yorkie-CA* (Fig. 8A,B). Furthermore, the myofibrils present in *yorkie-CA* at 32 h APF are less regularly organised compared to control, with actin showing less pronounced periodicity (Figure 6 supplement 1D). These ectopic localisation patterns of actin and myosin may explain muscle hyper-compaction at 32 h APF and the chaotic arrangement of myofibrils in *yorkie-CA* 90 h APF (see Figure 5C). Taken together, these data provide strong evidence that the Hippo pathway and its transcriptional regulator Yorkie contribute to the timing of myofibril assembly by regulating correct timing and levels of sarcomeric protein expression.

## Discussion

### The Hippo pathway and post-mitotic muscle fiber growth

Muscle fibers are often enormously large cells, which are densely packed with force producing contractile filaments and ATP producing mitochondria (Willingham et al., 2020). Thus, it appears logical that myofibril and mitochondria content primarily contribute to the size of an individual muscle fiber. To explore the link between myofibrillogenesis and muscle fiber growth, we used the largest *Drosophila* muscle cells, the indirect flight muscles, which are formed by fusion of several hundred myoblasts per muscle fiber and grow more than 10 times in volume after fusion within 72 hours. We provide strong genetic evidence that the regulation of Yorkie nuclear activity by the Hippo pathway is essential to allow post-mitotic flight muscle growth. Loss of function of Yorkie or one of the upstream STRIPAK complex members Slmap, Strip and Cka, as well as Dlg5, that regulate Hippo activity, all show the same phenotype: muscle atrophy during the post-mitotic muscle growth phase. Atrophy can be partially suppressed by over-expression of the apoptosis inhibitor DIAP1, which then results in small flight muscle fibers, emphasizing the essential role of Yorkie in promoting flight muscle growth.

How general is this role of Yorkie and the Hippo pathway in muscle fiber growth? Interestingly, muscle specific loss (*Dlg5-IR, Slmap-IR* and *yorkie-IR*) and gain of function of Yorkie (*hippo-IR, yorkie-CA*) results in viable but flightless animals. This suggests that the larval muscles and the other adult *Drosophila* muscle fibers such as leg and abdominal muscles can form and function normally in the absence of Yorkie and the Hippo pathway. One reason might be their slower growth rates and their more limited sizes compared to indirect flight muscles. A second reason may relate to the particular myofibrillogenesis mechanism in flight muscles. Flight muscles display individual distinct myofibrils, which all form simultaneously at about 32 h APF (Weitkunat et al., 2014) and then grow in length and diameter to match the volume increase of the muscle fiber (Spletter et al., 2018).

The closest mammalian homolog to insect flight muscles is the mammalian heart. Similar to flight muscles, the heart is a very stiff muscle using a stretch-modulated contraction mechanism (Shiels and White, 2008). After birth, mammalian cardiomyocytes stop dividing and all organ growth is achieved post-mitotically by size increase of the individual contractile cells. Interestingly, it was shown that mammalian YAP1 can promote cardiomyocyte survival and growth, as post-mitotic deletion of YAP1 results in increased fibrosis and cardiomyocyte apoptosis (Del Re et al., 2013). However, a role for the Hippo pathway in mammalian muscle is not limited to the heart. Constitutive expression of active YAP in adult mouse muscle fibers induces muscle fiber atrophy and deterioration of muscle function (Judson et al., 2013). Furthermore, it has been shown that the mammalian Hippo homolog Mst1 is a key regulator in fast skeletal muscle atrophy (Wei et al., 2013), and more importantly YAP promotes muscle fiber growth by its transcriptional activity requiring TEAD cofactors (Watt et al., 2015). This suggests that the Hippo pathway via its transcriptional regulators Yorkie/YAP/TAZ is a general regulator of muscle fiber growth and survival in animals.

### Yorkie targets

How does Yorkie mediate flight muscle growth? Our transcriptomics analysis revealed that mRNAs coding for sarcomeric and mitochondrial components are major transcriptional targets of Yorkie in flight muscle. In flies and mammals, Yorkie/YAP/TAZ require transcriptional cofactors of the Tead family (TEA domain containing) to bind to DNA. Interestingly, it has been shown that mammalian homologs Tead1-4 bind to a DNA motif called ‘muscle CAT’ (MCAT), a motif well-known to regulate cardiac skeletal sarcomeric protein expression, including cardiac Troponin T or cardiac actin (Carson et al., 1996; Farrance and Ordahl, 1996; Farrance et al., 1992; Wackerhage et al., 2014). The only *Drosophila* TEAD family member Scalloped binds to a very similar MCAT motif (Wu et al., 2008) and we found loss of *scalloped* shows the same phenotype as loss of function of *yorkie*. This strongly suggests that Yorkie and Scalloped cooperate in *Drosophila* muscle to transcriptionally boost sarcomeric gene expression to enable myofibril assembly and successively flight muscle fiber growth.

It has been shown that flight muscle fate is determined by the zinc finger transcriptional regulator Spalt major (Schönbauer et al., 2011). This includes the regulation of all flight muscle-specific sarcomeric components. How Yorkie cooperates with Spalt is to date an open question. One interesting link is that Yorkie and the Hippo pathway are required for the normal expression of *bruno1* (*bru1*, see Figure 7B). Bruno is the major splice regulator of flight muscle alternative splicing of sarcomeric proteins, downstream of Spalt (Spletter et al., 2015). Additionally, Bruno was shown to bind to 3’-UTRs of mRNA to regulate their translation efficiencies (Webster et al., 1997). Another RNA binding protein that requires the Hippo pathway for normal expression is Imp (IGF-II mRNA-binding protein, see Figure 7B). Imp has been shown to regulate the stability and translation of a number of F-actin regulators (Medioni et al., 2014), suggesting that post-transcriptional effects of Hippo signalling can play an important role, too. This may explain the strong up-regulation of Mhc and Act88F proteins levels in *yorkie-CA* muscle at 24 h APF despite little changes at the mRNA level. A similar cross talk between the Hippo pathway and translational regulation via the mTORC1 complex has been suggested in mammals (Hansen et al., 2015).

### Regulation of the Hippo pathway

How is the Hippo pathway regulated to induce sarcomeric protein expression at the correct time during muscle development? Salvador, Expanded, Merlin and Kibra, which regulate activity or plasma membrane localisation of Hippo in epithelial cells, all appear not to be required in flight muscles (Schnorrer et al., 2010)(and data not shown). However, we find that the core members of the STRIPAK complex Slmap, Strip and Cka as well as Dlg5 are required to regulate Hippo in muscle. Interestingly, Strip and Cka have been shown to bind to each other and regulate either Hippo or other signalling pathways in other *Drosophila* tissues including eye, testis or motor neurons (La Marca et al., 2019; Neal et al., 2020; Neisch et al., 2017). Slmap appears to be the specific adaptor to link STRIPAK to Hippo in muscle (our work) and epithelia in flies (Neal et al., 2020; Zheng et al., 2017) as well as in mammalian cell culture (Bae et al., 2017; Kwan et al., 2016). As Slmap has a single transmembrane domain, it is likely localised to the plasma membrane or t-tubules, membrane invaginations that form during myofibrillogenesis and link the plasma membrane to the forming sarcomeres (Razzaq et al., 2001). Slmap may either bind or recruit Dlg5 to the membrane, the latter was interestingly also found to bind to Mst1 (Hippo homolog) in mammalian cell culture (Kwan et al., 2016), which may boost the interaction of STRIPAK with Hippo, resulting in its effective de-phosphorylation. A detailed molecular model of Hippo signalling (Figure 9) remains speculative to date until the precise subcellular localisation of the key components, Hippo, Slmap and Dlg5 have been resolved in developing muscle.

**Figure 9.**
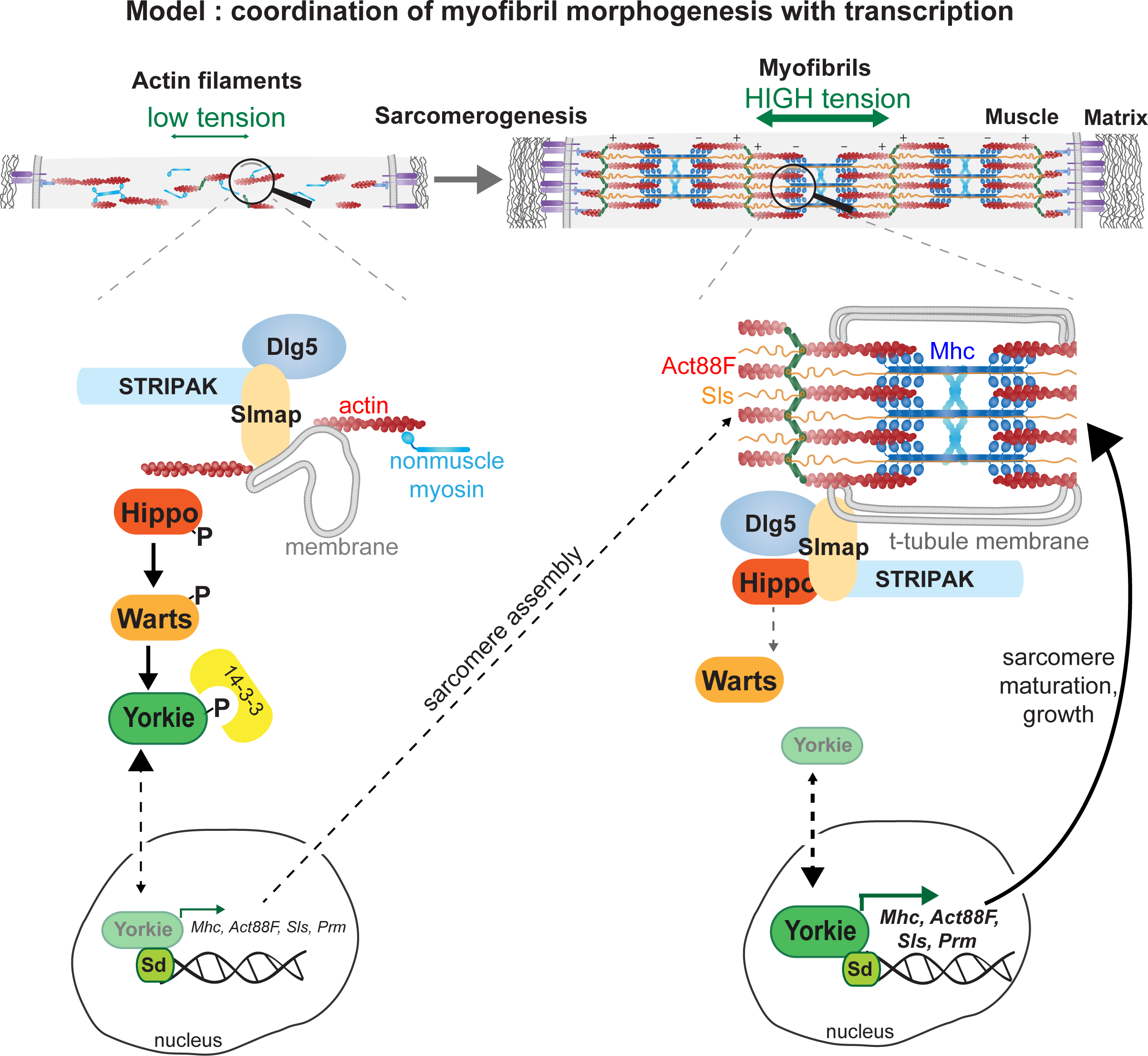
Model of Yorkie’s role how to coordinate myofibrillogenesis with transcription during muscle morphogenesis.

### A speculative model how myofibrillogenesis may coordinate with transcription

We have shown that mechanical tension in flight muscles increases during the early phases of myofiber development and identified it as a key regulator of myofibril assembly in flight muscle (Weitkunat et al., 2014). Concomitantly, transcription of sarcomeric components needs to be up-regulated to allow myofibril assembly and myofibril maturation. An attractive although speculative model places the Hippo pathway central to coordinate the assembly status of the myofibrils with the transcriptional status of the muscle fiber nuclei (Figure 9). In analogy to the epithelial cell cortex, we hypothesize that mechanical stretch produced by the actin cytoskeleton potentially via membranes, STRIPAK and Dlg5 may inhibit Hippo and thus promote the nuclear localisation of Yorkie, which in turn will boost transcription of mRNAs coding for sarcomeric components. This would be an analogous mechanism to the inhibition of Hippo by high tension produced by the cortical actomyosin and spectrin networks at the epithelial cortex resulting in epithelial tissue growth (Deng et al., 2015; Fletcher et al., 2015; 2018; Rauskolb et al., 2014). Interestingly, it was shown that filamentous actin (F-actin) levels can directly regulate Hippo pathway activity; more F-actin blocks Hippo signalling, resulting in epithelial tissue over-growth (Fernández et al., 2011; Sansores-Garcia et al., 2011; Yin et al., 2013). A similar feedback loop may enable the induction of more actin and myosin once the myofibrils have started to assemble and produce high levels of mechanical tension (Figure 9). This feedback model implies that only the successful completion of the myofibril assembly step then allows to fully boost sarcomeric mRNA transcription and translation to enable myofibril maturation and muscle fiber growth. This is consistent with the temporal separation of myofibril assembly and maturation in flight muscles (González-Morales et al., 2019; Shwartz et al., 2016; Spletter et al., 2018) Mechanical control of the Hippo pathway and YAP/TAZ localisation by the actomyosin cytoskeleton is also prominent in mammalian cells (Dupont et al., 2011; Wada et al., 2011). A similar mechanical feedback model as proposed here may therefore also be a relevant mechanism to coordinate post-mitotic cardiomyocyte and skeletal muscle growth or regeneration in mammals.

## Methods

### Fly strains and genetics

Fly strains were maintained using standard conditions. Unless otherwise stated, all experiments were performed at 27°C to improve RNAi efficiency. When applying temperature sensitive *Tub*-GAL80ts, fly crosses were kept at 18°C to suppress GAL4 activity and the white pre-pupae (0-30 min APF) were shifted to 31°C to allow GAL4 activity only at pupal stages. The pupae were then raised at 31°C until the desired age. Considering that pupae develop faster at 31°C compared to 27°C, timing was corrected by growing pupae at 31°C for 30 h age matching 32 h APF at 27°C, for 44 h matching 48 h APF and for 84 h matching 90 h APF at 27°C. The *Act88F*-GAL4 *UAS-Hippo* pupae were raised for 66 h at 18°C age matching 32 h APF at 27°C, and for 96 h at 18°C age matching 48 h APF at 27°C. RNAi stocks used were from the Vienna (Dietzl et al., 2007) or Harvard collections (Ni et al., 2011) and obtained from VDRC or Bloomington stock centers. All used fly strains are listed in Supplementary Table 4.

### Flight

Flight tests were performed as previously described (Schnorrer et al., 2010). One to three day old male flies were dropped into a 1 m long plexiglass cylinder with 8 cm diameter and 5 marked sections. Wild-type flies land in the upper 2 sections, whereas flies with defective flight muscles fall to the bottom of the tube immediately. For each genotype flight assays were performed two or three times with minimum 7 males each (for *yorkie-IR* females were used) (Supplementary Table 1).

### Pupal dissection and flight muscle stainings

For 24 h, 32 h and 48 h APF pupal dissections, the pupa was stabilized on a slide by sticking to double-sticky tape. The pupal case was removed with fine forceps. Using insect pins 2 or 3 holes were made in the abdomen to allow penetration of the fixative and pupae were fixed in 4% PFA (Paraformaldehyde) in PBST (PBS with 0.3% Triton-X) in black embryo glass dishes for 15 min at room temperature (RT). After one wash with PBST the pupae were immobilised using insect pins in a silicone dish filled with PBST and dissected similarly as described previously (Weitkunat and Schnorrer, 2014). Using fine scissors the ventral part of each pupa was removed, then the thorax was cut sagittally and the thorax halves were freed from the abdomen, leaving half thoraces with the flight muscles on them. These half thoraces were transferred to black embryo dishes and blocked for 30 min at room temperature (1/30 normal goat serum in PBST). F-actin was visualised with rhodamine-phalloidin (Molecular Probes, 1/500 in PBST) either alone or in combination with GFP-booster conjugated with Atto488 (ChromoTek, 1/200 in PBST) to visualise GFP fusion proteins. Half thoraces were incubated either 1 h at room temperature or overnight at 4°C. After 3 washes with PBST, half thoraces were mounted using Vectashield including DAPI (Biozol).

For 90 h APF pupal dissections, the head, wings, legs and abdomen were cut off the thorax with fine scissors, and the thoraxes were fixed for 20 min in 4% PFA in PBST at RT. After washing once with PBST, the thoraxes were placed on a slide with double-sticky tape and cut sagittally (dorsal to ventral) with a microtome blade (Pfm Medical Feather C35). These half thoraces were stained similarly to the early pupa half thoraces and mounted in Vectashield with DAPI using 2 spacer coverslips on each side.

### Image acquisition and processing

Image acquisition was performed with Zeiss LSM780 or LSM880-I-NLO confocal microscopes using Plan Neofluar 10x/0.30 NA air, Plan-Apo 40x/1.4 NA oil and Plan-Apo 63x/1.4 NA oil objectives. For all samples z-stacks were acquired. Image processing was done using Fiji (Schindelin et al., 2012). Digital cross-sections were created from z-stacks that covered the entire width of the flight muscle by drawing a straight line in the central part of the fiber and re-slicing. Fiber length and fiber cross-sectional area were measured with freehand drawing tools in Fiji based on phalloidin staining. To determine the average muscle fiber length per hemithorax, the length of all flight muscles for which both fiber ends were visible were measured and averaged.

### Myofibril length quantification and intensity profile plots

Myofibrils stained with phalloidin from a 40 μm × 20 μm × 2.5 μm confocal microscope stack were traced manually using Simple Neurite Tracer plug-in in Fiji (Longair et al., 2011). In each pupa, 10 myofibrils were traced and their average length was calculated. To visualise sarcomere periodicity, intensity profiles were plotted along a line based on actin labelling (phalloidin) using Fiji.

### Nuclei count

To count nuclei numbers of flight muscle fibers, half thoraces were stained with phalloidin (actin) and DAPI (nuclei) and imaged first using 10x objective to quantify the entire length of the fiber and then with 63x oil objective to visualize details. The acquired 63x z-stacks contain the entire muscle depth and about half of the muscle fiber length. Using Fiji’s multi-point tool all the nuclei in each z-stack were counted manually, using actin labelling as a landmark to visualize the borders of the fiber. Nuclei number for the entire fiber was calculated using the length of the entire fiber from the 10x image.

### Cryo cross-sections

Cryo cross-sections were performed as described previously (Spletter et al., 2018). Briefly, the pupal case was removed by fine forceps. Using insect pins 2 or 3 holes were made in the abdomen to allow penetration of the 4% PFA in PBST (PBS with 0.5% Triton-X) overnight at 4°C. Fixed pupae were sunk in 30% sucrose solution in PBST on a nutator at 4°C. Pupae were embedded in O.C.T. compound in plastic moulds (#4566, Sakura Finetek) and frozen on dry ice. Blocks were sectioned at 20 μm thickness on a microtome cryostat. Sections were adhered on glass slides coated with 1% gelatin, 0.44 mM chromium potassium sulfate dodecahydrate to improve tissue adherence. Sections on the slide were fixed for 1 minute in 4% PFA with PBS at room temperature, washed once for 5 minutes in PBS, incubated with rhodamine-phalloidin (1/500 in PBST) for 2 h at RT in a wet chamber, washed three times with PBST and mounted in Fluoroshield with DAPI.

### Quantifying GFP protein levels

GFP-tagged genomic fosmid fly lines (fTRG500 for Mhc-GFP and fTRG10028 for Act88F-GFP) (Sarov et al., 2016) were used for comparison of protein levels in *Mef2*-GAL4 driven knockdown of *yorkie* or gain of function *yorkie-CA* to *Mef2*-GAL4 control. Pupae of the different genotypes were dissected, stained and imaged on the same day in parallel under identical settings (master mixes for staining reagents, identical laser and scanner settings for imaging). Per hemi-thorax one or two different areas were imaged using 63x oil objective, zoom factor 2. In Fiji, five z planes at comparable positions in the muscles were selected for an average intensity projection of the volume into a 2D plane. In the 2D plane, two or three regions of 50 μm^2^ occupied by myofibrils (based on actin labelling) were selected. Mean intensities of each of these regions were averaged to calculate one value per hemi-thorax used for the quantification graphs. For each experimental day, the mean intensity of all wild type samples was set to 1 to calculate the relative intensity of the other genotypes. Data from a minimum of two independent experiments were plotted.

### RNA-isolation from developing flight muscle

For each replicate, flight muscles from seven *Mef2*-GAL4, *UAS-GFP-GMA* pupae at 24 h or 32 h APF were dissected in ice-cold PBS treated with DEPC using a fluorescent binocular. Flight muscles were collected in an Eppendorf tube and centrifuged at 2000 g for 5 min. The flight muscle pellet was re-suspended in TRIzol™, shock-frozen in liquid nitrogen and kept at −80°C.

RNA was isolated directly from the TRIzol muscle samples using a 96 well plate extraction kit (Direct-zol™-96 RNA, Zymo Research, #R2054): after thawing to room temperature in 1,5 ml Eppendorf tubes, the tissue samples were homogenized using a small pestle, followed by nucleic acid precipitation with 100% ethanol. This suspension was then transferred to the 96-well plate containing the purification columns. DNA digestion was performed ‘in column’ according to the kit instructions. Total RNA was eluted with 25 μl of RNase-free water and was quantified using the Quantifluor RNA System (Promega, #E3310).

### BRB-sequencing library preparation

RNA sequencing libraries were prepared using 20 ng of total RNA following the BRB-sequencing protocol (Alpern et al., 2019). Briefly, each RNA sample was reverse transcribed in a 96-well plate using SuperScriptTM II Reverse Transcriptase (Lifetech 18064014) with individual barcoded oligo-dT primers (Microsynth, Switzerland. For primer sequences see (Alpern et al., 2019)). Next, the samples were split into 3 pools, purified using the DNA Clean and Concentrator kit (Zymo Research #D4014), and treated with exonuclease I (New England BioLabs, NEB #M0293S). Double-stranded cDNA was generated by the second strand synthesis via the nick translation method. For that, a mix containing 2 μl of RNAse H (NEB, #M0297S), 1 μl of *E. coli* DNA ligase (NEB, #M0205 L), 5 μl of *E. coli* DNA Polymerase (NEB, #M0209 L), 1 μl of dNTP (0 .2 mM), 10 μl of 5x Second Strand Buffer (100 mM Tris, pH 6.9, AppliChem, #A3452); 25 mM MgCl_2_ (Sigma, #M2670); 450 mM KCl (AppliChem, #A2939); 0.8 mM β-NAD Sigma, N1511); 60 mM (NH_4_)_2_SO_4_ (Fisher Scientific Acros, #AC20587); and 11 μl of water was added to 20 μl of ExoI-treated first-strand reaction on ice. The reaction was incubated at 16 °C for 2.5 h. Full-length double-stranded cDNA was purified with 30 μl (0.6x) of AMPure XP magnetic beads (Beckman Coulter, #A63881) and eluted in 20 μl of water.

The Illumina compatible libraries were prepared by tagmentation of 5 ng of full-length double-stranded cDNA with 1 μl of in-house produced Tn5 enzyme (11 μM). After tagmentation the libraries were purified with DNA Clean and Concentrator kit (Zymo Research #D4014), eluted in 20 μl of water and PCR amplified using 25 μl NEB Next High-Fidelity 2x PCR Master Mix (NEB, #M0541 L), 2.5 μl of P5_BRB primer (5 μM, Microsynth), and 2.5 μl of Illumina index adapter (Idx7N5 5 μM, IDT) following program: incubation 72 °C—3 min, denaturation 98 °C—30 s; 15 cycles: 98 °C—10 s, 63 °C—30 s, 72 °C—30 s; final elongation at 72 °C—5 min. The fragments ranging 200– 1000 bp were size-selected using AMPure beads (Beckman Coulter, #A63881) (first round 0.5x beads, second 0.7x). The libraries were profiled with High Sensitivity NGS Fragment Analysis Kit (Advanced Analytical, #DNF-474) and measured with Qubit dsDNA HS Assay Kit (Invitrogen, #Q32851) prior to pooling and sequencing using the Illumina NextSeq 500 platform using a custom primer and the High Output v2 kit (75 cycles) (Illumina, #FC-404-2005). The library loading concentration was 2.2 pM and sequencing configuration as following: R1 6c / index 8c / R2 78c.

### Pre-processing of the data—de-multiplexing and alignment

The sample reads de-multiplexing was done using BRB-seqTools (http://github.com/DeplanckeLab/BRB-seqTools) as described before (Alpern et al., 2019). The sequencing reads were aligned to the Ensembl gene annotation of the *Drosophila melanogaster* BDGP6.23 genome using STAR (version 020201) (Dobin et al., 2013), and count matrices were generated with HTSeq (version 0.9.1) (Love et al., 2014).

### Bioinformatics analysis of BRB-Seq data

BRB-seq data quality was assessed in several ways. First, we excluded 5 samples from further analysis due to low numbers of aligned reads (< 500K; removed were 1x 24 h APF wild type, 1x *Dlg5-IR* 24 h, 1x *yorkie-CA* 24h and 2x 32 h APF wild type). Using raw read counts, we performed PCA analysis, calculated heatmaps and Pearson’s correlation in R (Version 3.3.1, https://cran.r-project.org/). One additional replicate from 32 h APF wild type was removed from further analysis as it represented a clear outlier. For the remaining samples, we performed library normalization (RLE) as well as differential expression analysis using DESeq2 (version 1.22.2, (Love et al., 2014)). Genes were considered differentially expressed with a log2FC ≥ |1| and an FDR ≤ 0.05. Functional enrichment analysis was performed on the differentially expressed genes using FlyEnrichr (Kuleshov et al., 2019). Data are available at the Gene Expression Omnibus database (Clough and Barrett, 2016), accession number GSE158957.

### Affinity Enrichment Mass Spectrometry

For each replicate, about 100 pupae staged from 24 h - 48 h APF of the genotypes *Mef2*-GAL4 (control) or *Mef2*-GAL4, *UAS-Dlg5-GFP* were collected and processed as described previously (Sarov et al., 2016). For each genotype, four replicates (100 pupae each) were snap-frozen in liquid nitrogen and ground to powder while frozen. The powder was processed as described (Hubner et al., 2010). Briefly, the cleared lysate was mixed with magnetic beads pre-coupled to a GFP antibody matrix to perform single step affinity enrichment and mass-spec analysis using an Orbitrap mass spectrometer (Thermo Fisher) and the QUBIC protocol (Hein et al., 2015). Raw data were analysed in MaxQuant version 1.4.3.22 (Cox and Mann, 2008) using the MaxLFQ algorithm for label-free quantification (Cox et al., 2014). The volcano plot was generated with Graphpad.

## Acknowledgements

The authors are indebted to Reinhard Fässler for hosting part of this work in his department, and to Jürg Müller and Anne-Kathrin Classen for hosting A.K.C during parts of this study. We are grateful to the IBDM and LIC (Albert-Ludwigs University of Freiburg) imaging facilities for help with image acquisition and maintenance of the microscopes, we acknowledge the France-BioImaging infrastructure supported by the French National Research Agency (ANR–10–INBS-04-01, Investments for the future). Fly stocks obtained from the Bloomington *Drosophila* Stock Center (NIH P40OD018537) and the Vienna *Drosophila* Resource Center (VDRC) were used in this study. The authors thank Pierre Mangeol and Clara Sidor for helpful discussions and critical comments for this manuscript.

## Figure legends

**Figure 1 supplement 1.**
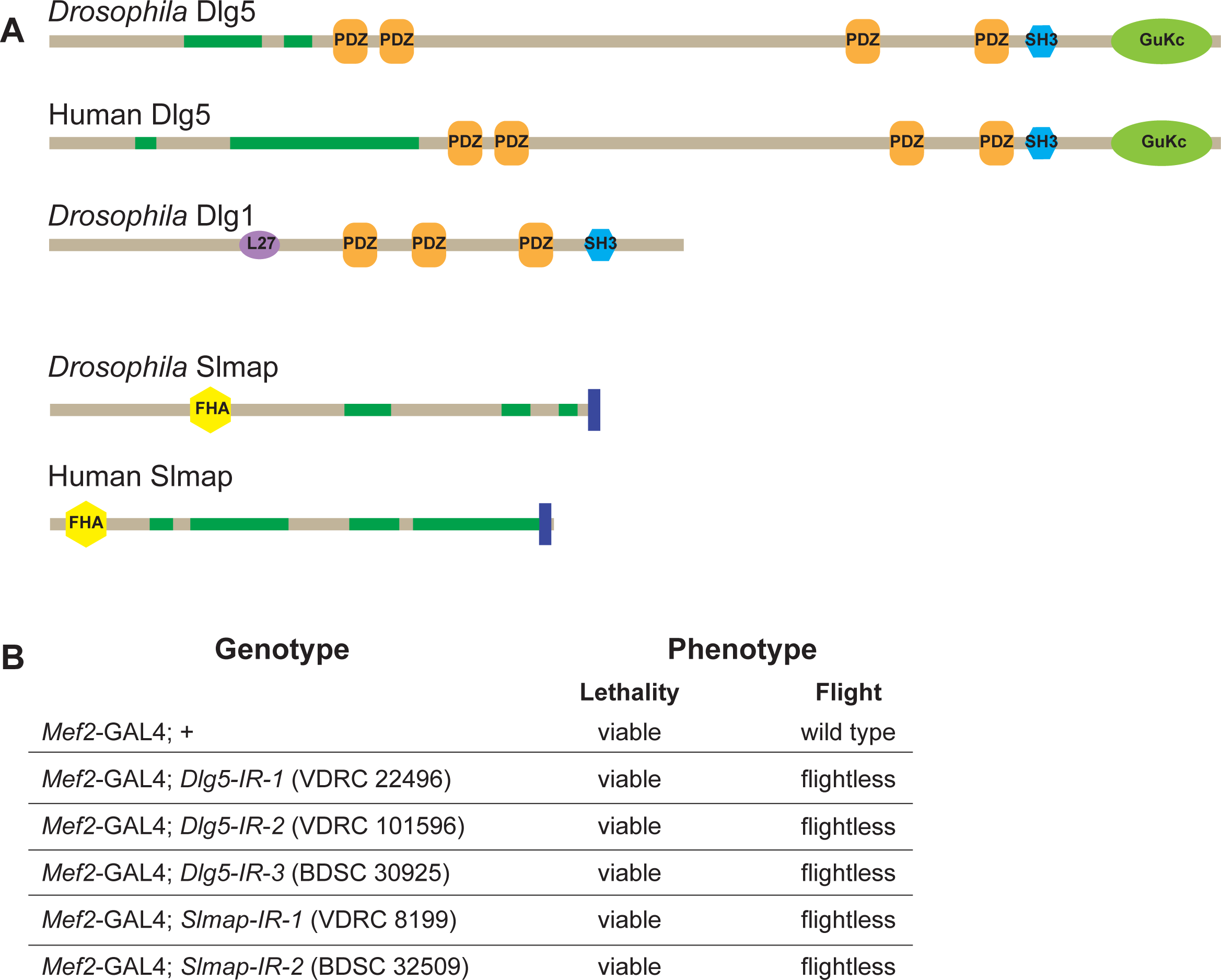
Dlg5 and Slmap proteins are conserved and required for flight. **A**. Protein domain organization comparing *Drosophila* Dlg5 to human Dlg5 and *Drosophila* Dlg1 (isoform M), as well as *Drosophila* Slmap to human Slmap. Note the conservation of Dlg5 and Slmap between *Drosophila* and human. PDZ: PSD95/Dlg1/ZO-1 domain; SH3: SRC Homology 3; L27 domain; GuKc: Guanylate kinase domain; FHA: Forkhead associated domain. **B**. Viability and flight tests of wild type (*Mef2*-GAL4 control) as well as *Dlg5* or *Slmap* knock-down genotypes. Note that knock-down for each of both genes results in viable but flightless adult flies.

**Figure 3 supplement 1.**
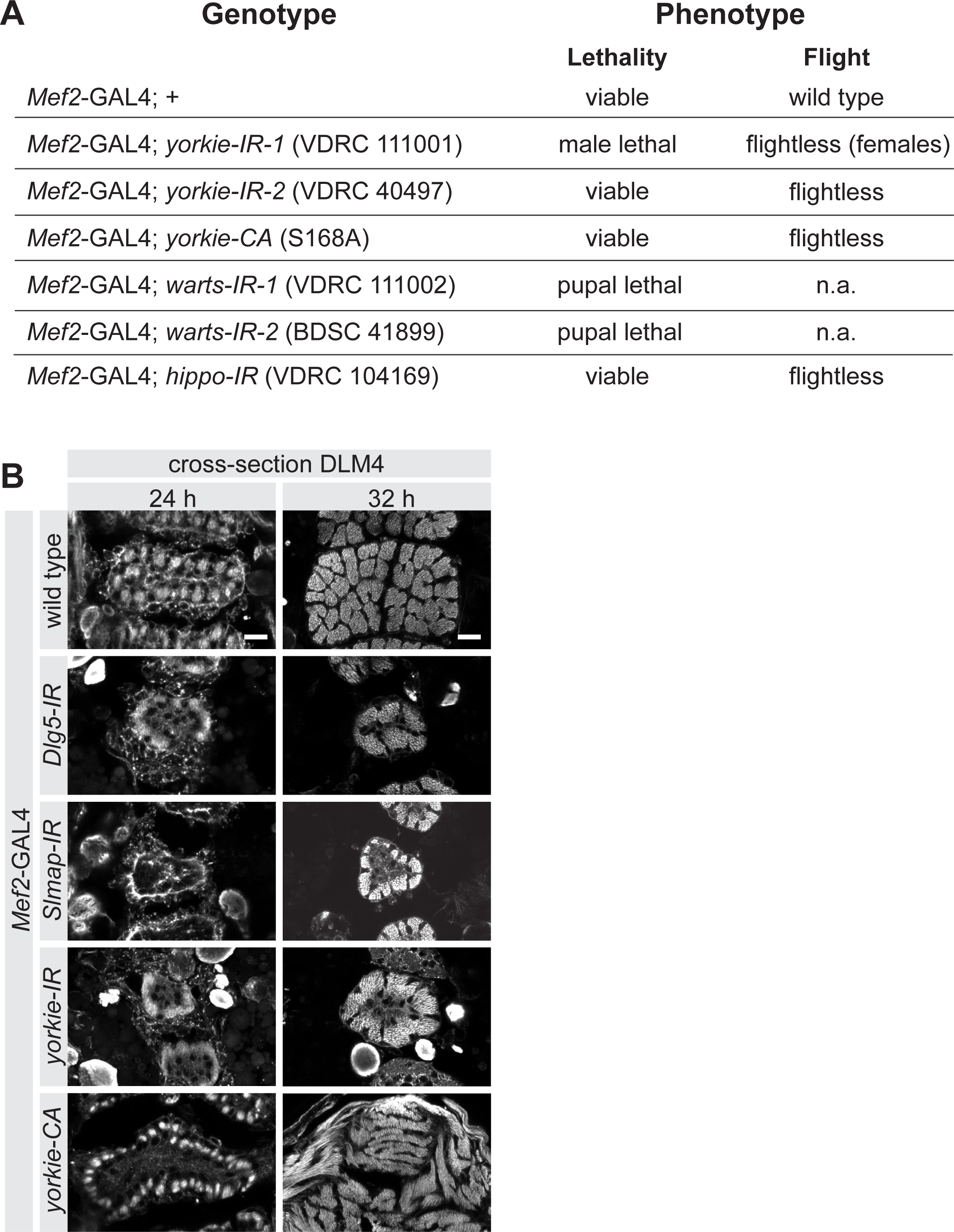
The Hippo pathway is required for muscle function. **A**. Viability and flight tests comparing wild type (*Mef2*-GAL4 control) to knockdown or over-expression of Hippo pathway components. **B**. Cryo cross-sections of dorsal longitudinal muscle 4 from 24 h and 32 h APF wild type, *Dlg5*, Slmap or *yorkie-IR* and *yorkie-CA* pupae. Scale bars represent 10 μm.

**Figure 4 supplement 1. The Hippo pathway is not required for myoblast fusion**

**A**. Quantification of nuclei number of DLM4 at 24 h APF from wild type, *Dlg5-IR, yorkie-IR* and *yorkie-CA Tub*GAL80ts *Mef2*-GAL4 (shifted to 31°C at 0 h APF). Fibers were stained for actin (phalloidin) and nuclei (DAPI). Images illustrating how the nuclei counting was done using longitudinal sections of DLM4. Since fiber widths vary in different genotypes, for illustration purposes different numbers of z-planes were maximum projected to result in comparable volumes. Dotted lines highlight the cell borders of the fibers. Scale bar represents 20 μm. **B**. Box plot showing total DLM4 nuclei numbers at 24h. Each dots represents one pupa. Student’s t test, n.s. p >0.05.

**Figure 6 supplement 1.**
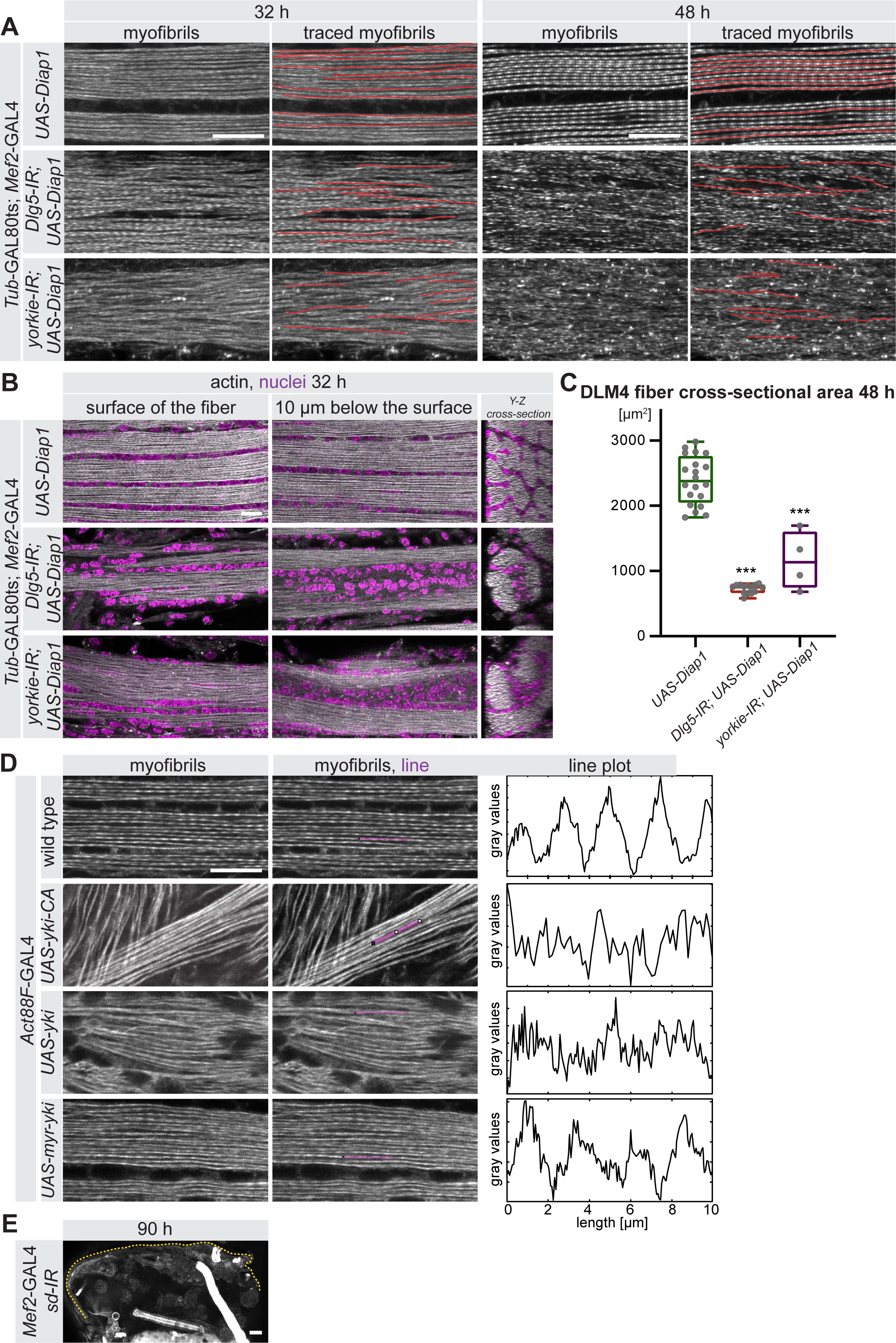
Myofibril tracing and nuclei positions in *Dlg5-IR* and *yorkie-IR* muscles. **A**. Myofibrils visualised by phalloidin from control, *Dlg5-IR* and *yorkie-IR UAS-Diap1 Tub*GAL80ts *Mef2*-GAL4 muscles at 32 h and 48 h APF (shifted to 31°C at 0 h APF). Myofibrils were traced with Simple Neurite Tracer and traces are highlighted in red. Note that in *Dlg5-IR* and *yorkie-IR* myofibrils traces are short. Scale bars represent 10 μm. **B**. Flight muscles stained for actin (phalloidin) and nuclei (DAPI) from control, *Dlg5-IR* and *yorkie-IR UAS-Diap1 Tub*GAL80ts *Mef2*-GAL4 muscles at 32 h (shifted to 31°C at 0 h APF). Note that nuclei fail to distribute between the myofibril bundles but cluster centrally in *Dlg5-IR* and *yorkie-IR* fibers. Scale bar represents 10 μm. **C**. Box plot of myofiber cross-sectional areas from cryo cross-sections of DLM4 from control, *Dlg5-IR* and *yorkie-IR UAS-Diap1 Tub*GAL80ts *Mef2*-GAL4 muscles at 48 h APF. Student’s t test, *** p-value <0.001. **D**. Intensity profiles of control myofibrils compared to UAS-*yorkie-CA*, UAS-wild-type-*yorkie* or UAS-*myr-yorkie* expressed with the post-mitotic *Act88F*-GAL4 driver at 32 h APF. Note the less pronounced actin periodicity in *UAS-yki* and *UAS-yki-CA* compared to wild-type control. Scale bars represent 10 μm. **E**. 90 h APF half thorax from *scalloped-IR Mef2*-GAL4. The dotted lines highlight the cuticle. Scale bar represents 50 μm.

**Figure 7 supplement 1.**
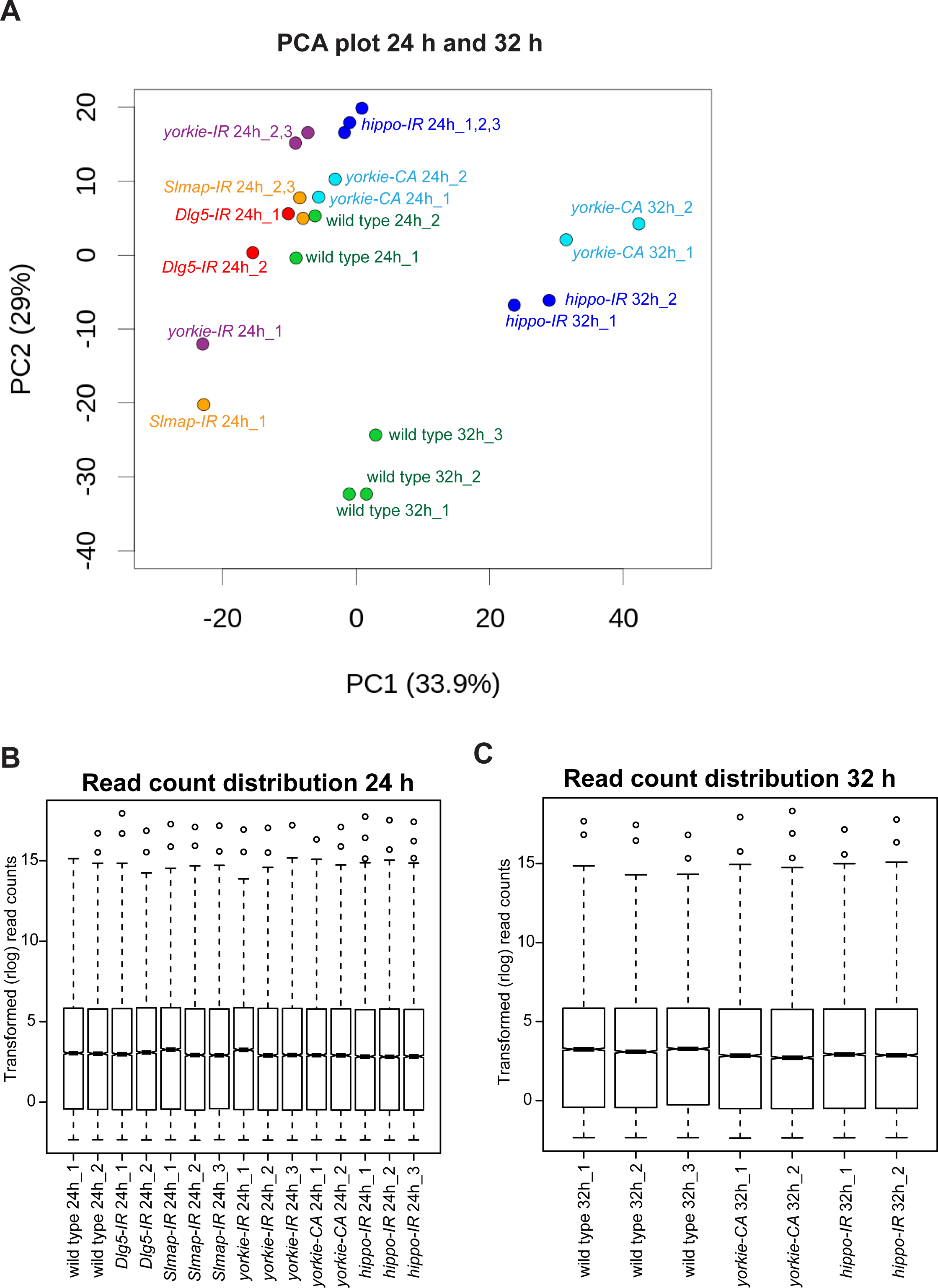
BRB sequencing read counts and PCA analysis. **A**. Principle component analysis (PCA) of BRB-sequencing replicates. Note the distinct clustering of 24 h from 32 h APF samples. Most but not all samples of similar genotypes cluster together. B. Transformed read count distributions of the 24 h and 32 h APF samples.

**Supplementary Table 1**

Data table containing data from Figures 1, 2, 3, 4, 5, 6 and 8.

**Supplementary Table 2**

Table listing the expression levels and fold changes as well as normalised p-values of the BRB-SEQ data compared to the wild-type controls. All or the only the significantly different genes are listed.

**Supplementary Table 3**

GO enrichment terms of the significantly different gene lists of the various genotypes and time points.

**Supplementary Table 4**

Table listing all *Drosophila* strains and major reagents used in the study

